# Disruption of genes associated with Charcot-Marie-Tooth type 2 lead to common behavioural, cellular and molecular defects in *Caenorhabditis elegans*

**DOI:** 10.1101/605584

**Authors:** Ming S. Soh, Xinran Cheng, Jie Liu, Brent Neumann

## Abstract

Charcot-Marie-Tooth (CMT) disease is an inherited peripheral motor and sensory neuropathy. The disease is divided into demyelinating (CMT1) and axonal (CMT2) neuropathies, and although we have gained molecular information into the details of CMT1 pathology, much less is known about CMT2. Due to its clinical and genetic heterogeneity, coupled with a lack of animal models, common underlying mechanisms remain elusive. In order to understand the biological importance of CMT2-casuative genes, we have studied the behavioural, cellular and molecular consequences of mutating nine different genes associated with CMT2 in the nematode *Caenorhabditis elegans* (*lin-41/TRIM2, dyn-1/DMN2, unc-116/KIF5A, fzo-1/MFN2, osm-9/TRPV4, cua-1/ATP7A, hsp-25/HSPB1, hint-1/HINT1, nep-2/MME*). We show that *C. elegans* defective for these genes display debilitated movement in crawling and swimming assays. Severe morphological defects in cholinergic motors neurons are also evident in two of the mutants (*dyn-1* and *unc-116*). Furthermore, we establish novel methods for quantifying muscle morphology and use these to demonstrate striking loss of muscle structure across the mutants that correspond with reductions in muscle function. Finally, using electrophysiological recordings of neuromuscular junction (NMJ) activity, we uncover reductions in spontaneous postsynaptic current frequency in *lin-41, dyn-1, unc-116* and *fzo-1* mutants. By comparing the consequences of mutating numerous CMT2-related genes, this study reveals common deficits in muscle structure and function, as well as NMJ signalling when these genes are disrupted.

## Background

First identified in 1886, Charcot-Marie-Tooth (CMT) disease is the most common inherited peripheral neuropathy with an estimated prevalence of 1 in 2,500 individuals (Barisic et al., 2007, Barreto et al., 2016). Common clinical features of CMT include slowly progressive distal muscle weakness and atrophy, distal sensory impairment, foot deformities, secondary steppage gait and mobility impairment (Barisic et al., 2007, Reilly et al., 2011). There is currently no cure for CMT, and patients frequently suffer lifelong disabilities. While CMT can be inherited in an autosomal dominant, autosomal recessive or X-linked manner, it is predominantly (∼90% of cases) inherited autosomal-dominantly (d’Ydewalle et al., 2012). The disease is typically divided into two main categories: type 1 CMT (CMT1), also known as demyelinating neuropathy, affects the myelin sheath surrounding the axon resulting in decreased nerve conduction velocities (less than 38 m/s); whereas axonal Type 2 CMT (CMT2) affects the nerve axon and normally does not affect the speed of nerve conduction (Barisic et al., 2007, d’Ydewalle et al., 2012, Shy and Patzkó, 2011).

CMT subtype 2A (CMT2A) is the most prevalent among CMT2 and accounts for approximately 20% of CMT2 cases (Cartoni and Martinou, 2009). Currently, more than 20 genetic loci have been associated with CMT2, with each causative gene encoding a different protein(Bird, 1993-2018). These proteins are associated with an eclectic mix of cellular functions. For instance, the causative gene for CMT2A, *MFN2*, encodes mitofusin 2, a GTPase that regulates outer mitochondrial membrane fusion (Chandhok et al., 2018, Züchner et al., 2004). Conversely, *HSPB1*, the gene involved in CMT2F, encodes a heat shock protein with molecular chaperone functions (Ackerley S et al., 2006). Other causative genes are involved in protein translation (*GARS, AARS MARS, HARS*), osmotic regulation (*TRPV4*), signalling pathways and cell adhesion (*LRSAM1*), and neuroprotection (*TRIM2*). To further complicate matters, advancements in genetic screening has led to the discovery of additional CMT2 causal genes, with the latest one, *ATP1A1*, encoding a sodium/potassium-transporting ATPase, identified in 2018 (Lassuthova et al.). This results in a constant modification of the disease classification system in order to include the newer members, making the nomenclature imprecise and confusing (Patzkó and Shy, 2012). Besides causal genes, the age and mechanisms of onset, disease progression and severity also vary from one CMT2 subtype to another, and from patient to patient within the same subtype (Bird, 1993-2018). Due to its genetic and clinical heterogeneity coupled with the relative neoteric discovery of causal genes, it remains to be determined how mutations in a vast range of proteins with distinct functions all lead to CMT2. This has hindered the development of therapeutics for the disease. To aid our understanding of CMT2 pathophysiology, development of animal models which carry the mutant causal genes has played a crucial role. Unfortunately, unlike CMT1, the scarcity of CMT2 animal models has caused the study of underlying molecular mechanism of the axonal neuropathy to trail behind its demyelinating counterpart (Cartoni and Martinou, 2009).

*C. elegans* has emerged as one of the most widely used animal model systems to address questions regarding cellular and molecular aspects of human disease *in vivo*. It is also one of the few available animal models that are suitable for conducting cost-effective, rapid drug screening due to its ease of culturing, small body size (∼1 mm in length), high brood size (∼300 offspring), and short generation time (∼3 days to adulthood) (Markaki and Tavernarakis, 2010). Furthermore, its conserved musculature features, transparent body and ease of genetic manipulation make *C. elegans* a highly desirable model organism to study cellular morphology and human diseases (Laranjeiro et al., 2017). Approximately 60-80% of human genes have an orthologue in the *C. elegans* genome, and 42% of these genes are disease-related (Markaki and Tavernarakis, 2010). While *C. elegans* is an established model to study other pathologically similar neurodegenerative diseases such as amyotrophic lateral sclerosis, only a handful of the many CMT subtypes have been studied in the nematode.

To advance our understanding of the function of genes associated with CMT2, we have assessed the cellular and behavioural consequences of loss-of-function or null mutations in nine orthologous causative genes in *C. elegans*. More specifically, we studied crawling and swimming behaviour, both individually and within populations, examined the morphology and function of cholinergic motor neurons and body wall muscles, and analyzed neuromuscular junction transmission in these animals. As first described in this study, we have also devised methodology for quantitatively measuring muscle cell area and the length of muscle filaments in order to non-subjectively determine and compare the extent of muscle defects. Our study reveals common deficiencies in muscle structure and function, and consequent locomotion difficulties amongst our mutants. Strikingly, our study also reveals shared deficits in cholinergic neurons, as well as a prevailing site of dysfunction at the neuromuscular junction in the genes associated with the most severe movement and muscle phenotypes. Using *C. elegans* for comprehensive analyses of the orthologous genes causing nine different CMT2 subtypes, our study identifies common and diverse sites of defects, reveals novel insights into the functions of the CMT2-related genes, and paves the way for future studies aimed at generating effective therapeutics.

## Results

### Individual CMT2-associated mutants exhibit movement defects similar to CMT2 patients

The genes associated with CMT2 investigated in this study, along with their encoded proteins, are listed in Table 1 and Table S1. In addition to the genetic and clinical heterogeneity, our understanding of CMT2 is further complicated by numerous different alleles of the same gene being associated with each subtype. For example, more than 100 different mutations in the *MFN2* gene have been identified in CMT2A patients (Iapadre et al., 2018). Moreover, these mutations affect the encoded mitofusin 2 protein in different ways: some have no effect on protein function, some are gain-of-function, and others are loss-of-function that block its function in mitochondrial fusion (Detmer and Chan, 2007, El Fissi et al., 2018). To counteract some of these issues, instead of generating strains carrying individual patient-specific mutations, we have assessed the consequences of genetic null or loss-of-function alleles in order to expand our global understanding of the role of genes implicated in CMT2. We first sought to test whether dysfunction of these genes in *C. elegans* would result in similar behavioural symptoms as their human counterparts. As defective locomotion is a major symptom in CMT2 patients, we explored the main two means of locomotion displayed by *C. elegans*: swimming in liquid and crawling on solid media. Both of these activities involve molecular mechanisms conserved with humans, including depletion of energy, muscle fatigue and increased mitochondrial oxidation in muscle cells (Laranjeiro et al., 2017).

**Table 1.**
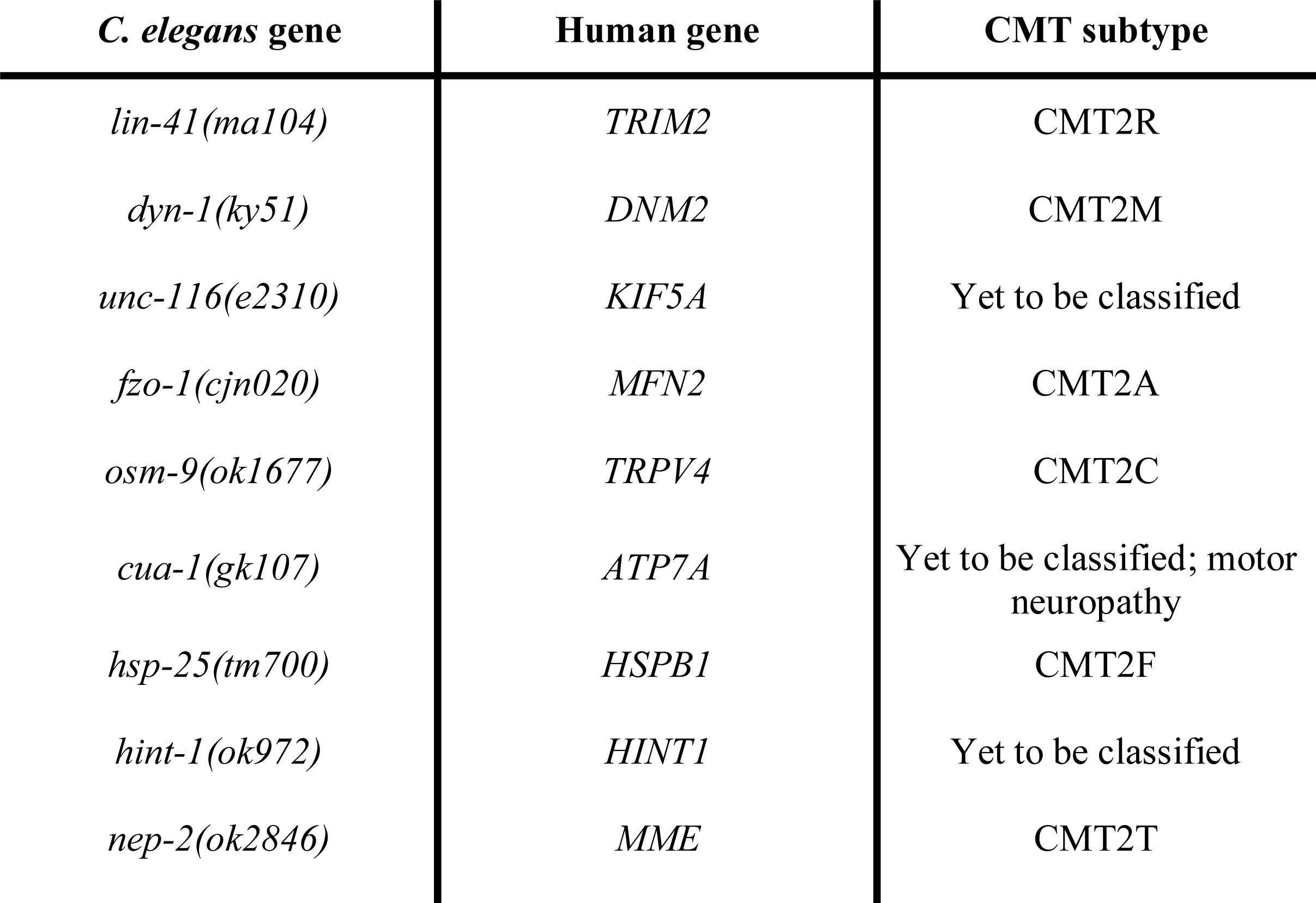
List of genes associated with CMT2 explored in this study.

To assess swimming behaviour, we performed thrash assays. We placed a single worm in liquid and counted the number of thrashes per minute, defined as a complete cycle of maximum bend on one side of the body (Fig. 1A). We compared the thrash rate between wild-type animals and those carrying mutations in the CMT2-related genes in four synchronized ages: the first larval stage 1 (L1), the final larval stage 4 (L4), 3-day old adult (A3) and 7-day old adult (A7). Mutations in seven of the nine CMT2-associated genes led to progressive deteriorations in locomotion, especially in the older animals (Fig. 1B and S1A). Interestingly, the onset of the defect varied between different genetic mutations. Worms carrying mutations in *dyn-1, unc-116* or *cua-1* presented lower thrash rates across all ages tested compared to wild-type, whereas the swimming ability of *lin-41(ma104)* and *osm-9(ok1677)* mutants were only significantly reduced in adult animals. Similarly, adult worms lacking functional FZO-1 also displayed significantly decreased thrash rates. However, the younger L4 worms did not share that phenotype, even though L1 *fzo-1(cjn020)* worms were found to be defective. The strictly age-dependent trend was also absent in *hsp-25(tm700)* animals, as only the L4 and A7 worms had significantly lower thrash rates. Surprisingly, mutation of *hint-1* (unclassified CMT) or *nep-2* (CMT2T) did not reduce the thrash rates at any age tested. The *hint-1(ok972)* allele deletes the entire *hint-1* gene, while the *nep-2(ok2846)* allele is a 362 bp deletion across the second and third exons of *nep-2*. Thus, although we expect both to strongly affect gene function, our analyses suggest that neither *hint-1* nor *nep-2* is important for the normal swimming behaviour of *C. elegans*.

**Figure 1.**
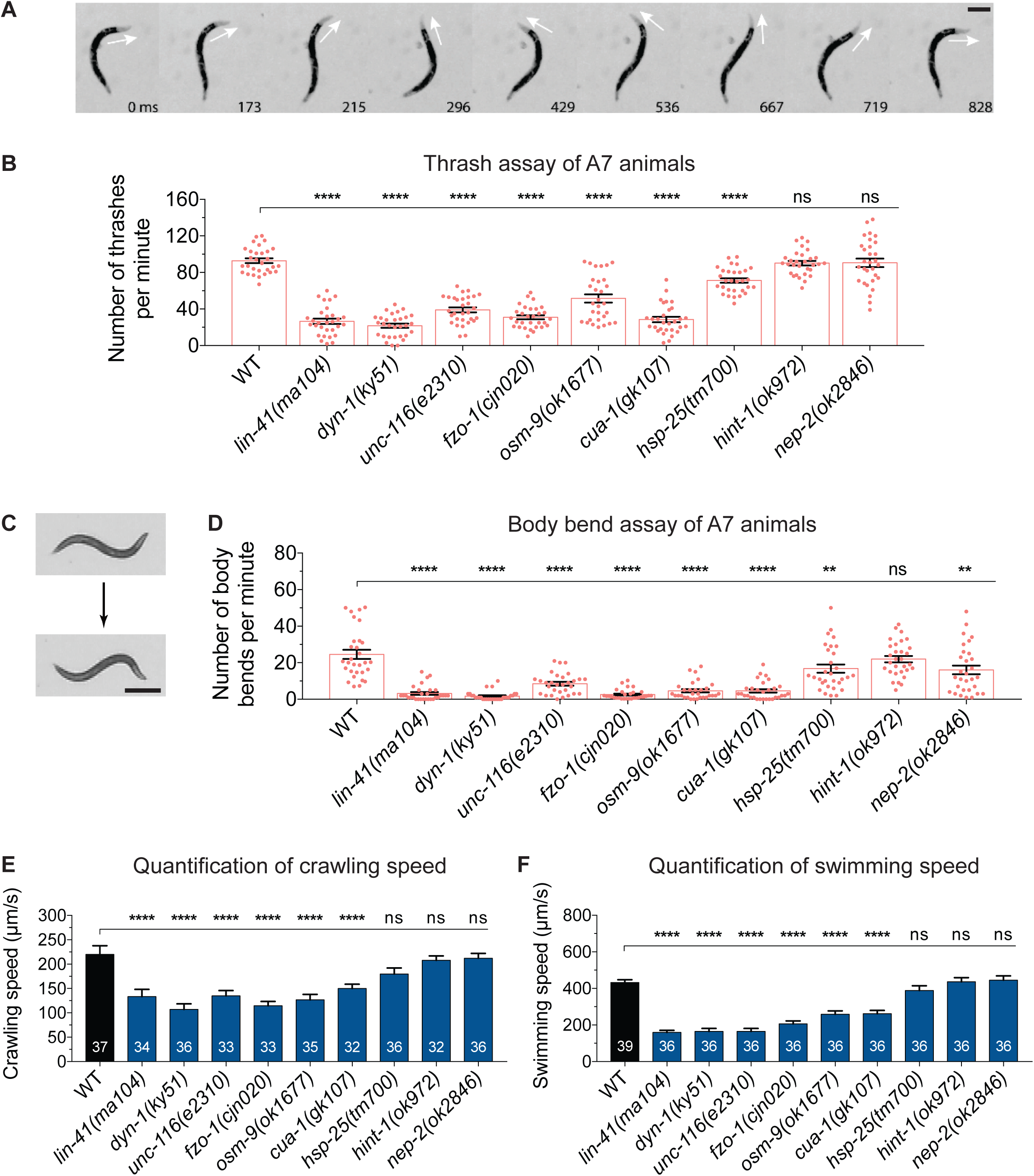
Motility assays in *C. elegans* mutants. (**A**) An animal performing a single thrash in liquid culture. The head begins facing right, moves to the left and then returns to the right. Arrows points to direction of the head; time in milliseconds displayed at bottom relative to the first image. Scale bar represents 250 µm. (**B**) Number of thrashes per minute calculated for 7-day old adult (A7) N2 wild-type (WT) and the nine mutant strains shown. See Figure S1A for thrash assays performed across different ages. (**C**) An animal undergoing one complete body bend on nematode growth media (NGM) agar. One body bend is defined as the maximum bending of the section behind the pharynx in the opposite direction, from top to bottom picture. Scale bar represents 500 µm. (**D**) Quantification of number of body bends of WT and CMT2 mutant A7 animals crawling on solid NGM agar. Number of body bends across different ages is included in Figure S1B. Each dot in (B) and (D) represents a single animal (*n* ≥ 30). (**E**) The rates of crawling on solid NGM agar and, (**F**) swimming in M9 liquid for 3-day old adult WT and nine CMT2 mutant worms. The swimming and crawling velocity were calculated using WormLab software; *n* values are within each bar. One-way ANOVA with Dunnett’s post hoc tests were used to compare swimming and crawling behaviour between WT and mutant animals in (B), (D), (E) and (F). Data is represented as mean ± S.E.M. \*\**P* < 0.01, \*\*\**P* < 0.001, \*\*\*\**P* < 0.0001, *ns* = not significant.

While swimming is a more energy-demanding exercise, worms crawling on a solid surface face far greater mechanical resistance than those swimming in liquid (Laranjeiro et al., 2017). To quantify crawling capacity, we manually counted the number of body bends of individual worms crawling on NGM agar over a three-minute time frame (Fig. 1C, D). Similar to the swimming deficiencies, all but one of the mutant strains displayed a considerably lower number of body bends over time compared to wild-type (Fig. 1D and S1B). Mutation of *dyn-1* or *fzo-1*, the causal genes for CMT2M and CMT2A respectively, led to the largest reductions in crawling rate, with younger worms presenting 60-80% reductions and older worms registering an average of just 2 body bends per minute compared to 25 in the wild-type. Unlike thrashing, the crawling ability of *lin-41(ma104)* mutants was impaired from birth, and drastically worsened in adulthood. In contrast, while *unc-116(e2310), osm-9(ok1677)* and *cua-1(gk107)* worms experienced significantly lower body bending rates across all ages, the decline was more gradual. Mutations in both *hint-1* and *nep-2* again had minimal effects on animal movement, although a significant reduction was observed in *nep-2(ok2846)* mutants at the oldest age analyzed.

Next, we quantified swimming and crawling speeds using an automated worm tracking program (WormLab), which we used to simultaneously track and analyze multiple worms. We focused on A3 worms as the locomotion defect of most CMT2 mutants became prominent at this age. Similar to the thrash and body bend assays, the majority of CMT2 mutants displayed at least 30-40% declines in swimming and crawling speeds compared to wild-type worms (Fig. 1E, F). Overall, these data establish that *C. elegans* carrying mutations in CMT2-associated genes experience similar movement defects to their human counterparts. Moreover, it separates the mutations into three categories in the worm: severe (*lin-41, dyn-1, unc-116, fzo-1*), intermediate (*osm-9, cua-1, hsp-25*), and mild *(hint-1, nep-2*).

### Populations of CMT2-associated mutants experience declines in motility over time

We next investigated whether locomotor defects were also observable on a population level. We recorded the motility of 30-50 worms from each group for three hours using an automated movement detection instrument (WMicrotracker). The system works by detecting the interference of infrared microbeams as a result of animal movement, whereby motile worms disrupt the infrared beam more frequently than stationary worms (Simonetta and Golombek, 2007). Beam disruption is registered and converted into an activity count. To obtain consistently reproducible data, we optimized methods to generate synchronized populations of worms (Fig. S2). To obtain large numbers of synchronized, gravid adults were first bleached to release the eggs, which were incubated overnight for hatching (Solis and Petrascheck, 2011). Hatched L1 worms were then plated the next day and left to grow until they reached adulthood. Previous studies have used fluorodeoxyuridine (FUdR) as an inhibitor of DNA synthesis to prevent adult worms from reproducing. However, since FUdR can affect *C. elegans* stress responses and lifespan (Van Raamsdonk and Hekimi, 2011, Feldman et al., 2014), as well as eliciting responses after metabolism in *E. coli* (the food source for *C. elegans*) (García-González et al., 2017), we instead manually separated adult worms from their progeny until they reached the 3-day old adult stage for WMicrotracker experiments. The separation was achieved by washing the plate containing adult worms and progenies into a tube, and letting the adult worms precipitate by gravity (Fig. S2). The synchronized A3 worms were then plated into a 96-well plate (at least 6 wells per group per experiment) and movement measured with the WMicrotracker instrument. During this optimization, we found that having 30-50 worms per well was optimal to prevent overcrowding, which we observed to inversely affect motility.

From the WMicrotracker screens, the wild-type populations of worms started with a motility count of about 420 arbitrary units (au), but slowly became less motile, reaching a plateau of around 330 au over the final hour of analysis (Fig. 2A-C). The average motility of wild-type worms during the 3-hour experiments was 365 au (Fig. 2D). In stark contrast, both *dyn-1(ky51)* and *unc-116(e2310)* mutants consistently recorded motility counts below 200 au (Fig. 2A, B, D). Populations of *lin-41(ma104), fzo-1(cjn020)* and *cua-1(gk107)* animals also experienced significantly reduced movement, registering between 220 and 330 au throughout the experiments (Fig. 2A, B, D). Worms lacking *hsp-25* had a slight (∼12%), but statistically significant reduction in motility, whereas mutation of *osm-9, hint-1* or *nep-2* had no effect on movement (Fig. 2A-D). In summary, these population-based motility assays revealed comparable movement defects to what was observed with individual worms in our previous locomotion assays. These automated, time-effective WMicrotracker assays could prove highly advantageous for drug screening approaches to identify compounds that are able to reverse the movement defects in these animals.

**Figure 2.**
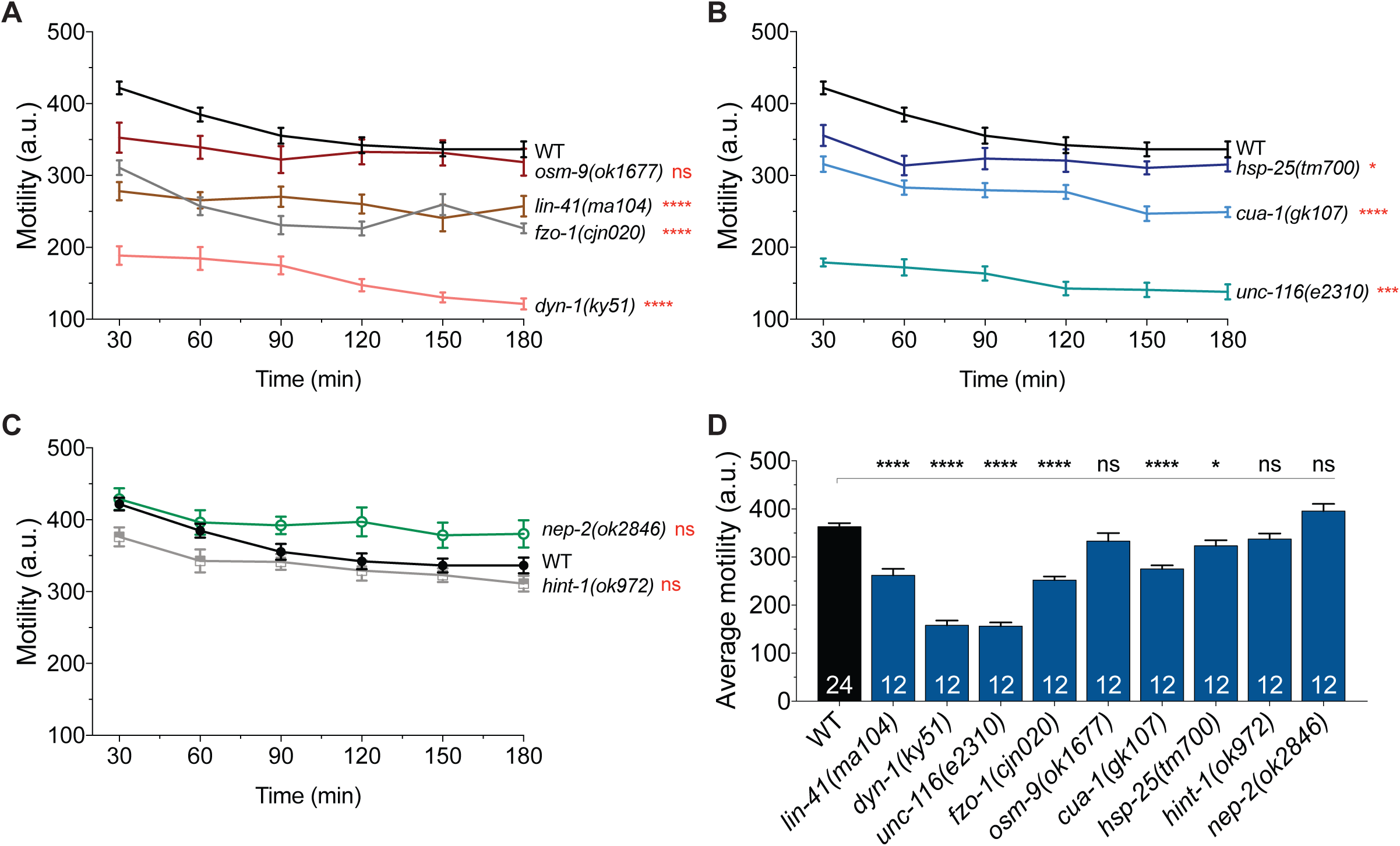
High-throughput analysis of swimming behaviour in animal populations. (**A-C**) Activity count graphs from WMicrotracker experiments. Motility of swimming animals (30-50 animals per well, 6 wells per genotype per experiment) was measured over a 3 h period in duplicate runs (*n* = 12 wells). Locomotor activity was compared between wild-type (WT) animals and the mutant strains indicated. (**D**) Summary and comparison of the average activity count for individual genotypes. The number of wells analyzed is reported in each bar. Data is represented as mean ± S.E.M. \**P* < 0.05, \*\*\*\**P* < 0.0001, *ns* = not significant from one-way ANOVA with Dunnett’s post hoc tests; *a.u.* arbitrary units.

### Mutation of CMT2-related genes results in defective cholinergic motor neuron morphology

Degeneration of peripheral axons of motor neurons is one of the key features of CMT2. In mammals, these neurons are cholinergic in nature and extend axons from the spinal cord to the periphery where they innervate skeletal muscle. In *C. elegans*, there are 17 classes of cholinergic motor neurons, where four of these, VA, VB, DA and DB, are known to innervate body wall muscles that are essential for locomotion (Altun and Hall, 2009b, White et al., 1986). In this study, we focused on the axonal morphology of *C. elegans* cholinergic motor neurons, DA and DB, which innervate dorsal wall muscles. Unfortunately, we could not accurately identify VA and VB neurons due to their overlapping positions with the other cholinergic neurons on the ventral side of the worm. We examined the commissures of DA and DB neurons in laterally-positioned A3 stage wild-type and CMT2 mutant worms using a cholinergic-specific, green fluorescent protein (GFP)-expressing transgene [*vsIs48(Punc-17::GFP)*] (Fig. 3A). Worms were qualified as defective or not based on the presence of one or more specific defects in the DA or DB neurons: absent commissure (missing), axonal guidance defect (guidance), visible breakage along the commissure (break), and/or short neurite extending less than half the body width (short) (Fig. 3B).

**Figure 3.**
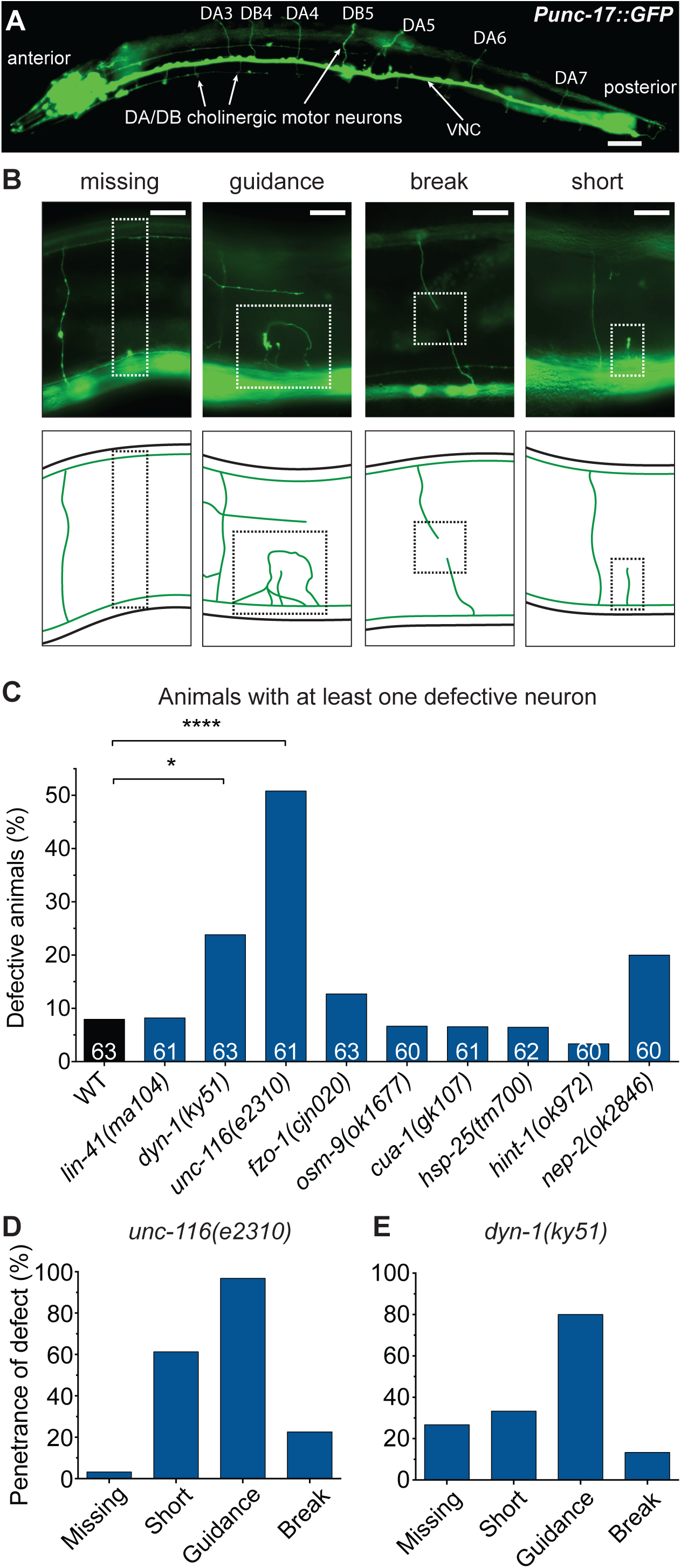
Morphological analysis of cholinergic motor neurons. (**A**) Full body image of a 3-day old adult animal carrying the *vsIs48*(*Punc-17::gfp*) transgene, which produces GFP expression in all cholinergic neurons. VNC = ventral nerve cord. (**B**) Images (top) and schematics (bottom) of specific morphological phenotypes: absent commissure (missing), wayward commissure that failed to connect with the dorsal nerve cord (guidance), a gap or break within a commissure (break), and short neurite (short). The phenotypes are highlighted with the dashed boxes. (**C**) Quantification of morphological defects in WT and CMT2-associated mutant animals. Animals were scored visually for defects. Bar represents percentage of defective animals; *n* values are shown in each bar. Chi-square test with false discovery rate were performed to compare defects between WT and mutant animals. \**P* < 0.05, \*\*\*\**P* < 0.0001. (**D**) Proportion of specific morphological defects in *unc-116(e2310)*, and (**E**) *dyn-1(ky51)* animals. Scale bar represents 40 µm in (A) and 8 µm in (B).

Of the nine CMT2-related genes, mutation of *unc-116* caused the most severe axonal defects in DA and DB motor neurons, with half of the worms displaying defects (Fig. 3C). Nearly all of these defective worms had misguided commissures, and more than half had stunted neurite growth (Fig. 3D). This trend was also observed in *dyn-1* mutants, the second most defective amongst the nine genes, with guidance and short commissure defects emerging again as the most common types of architectural defects (80% and 33% respectively; Fig. 3E). While mutation of *nep-2* saw an increase in DA/DB defect to 20%, this was not statistically significant (*P* > 0.05) (Fig. 3C). Worms carrying mutations in the remaining six genes did not exhibit substantial morphological defects in DA and DB neurons, suggesting that neuronal degeneration may not be a major driver of the movement defects in these animals. However, in addition to the 4 classes of cholinergic motor neurons, the *C. elegans* body wall muscles are also innervated by GABAergic motor neurons not present in mammals, but whose degeneration may play a role in the conspicuous locomotion impairments.

### CMT2-associated mutants display degeneration of body wall muscles

Due to axonopathy, the skeletal muscles of CMT2 patients gradually undergo denervation, leading to muscle atrophy and weakness (Bird, 1993-2018). Hence, we next sought to characterize the morphology of muscle cells in our nematode models. *C. elegans* possesses 95 obliquely striated body wall muscle cells, equivalent to mammalian skeletal muscles, that are arranged as staggered pairs in four longitudinal quadrants along the length of the worm (Fig. 4A, B) (Altun and Hall, 2009a, Gieseler et al., 2005). Individual muscle cells are connected intercellularly through gap junctions, allowing electrical signals to be transmitted rapidly between cells. Like motor neurons, functional body wall muscles are essential for *C. elegans* locomotion, including crawling and swimming (Altun and Hall, 2009a).

**Figure 4.**
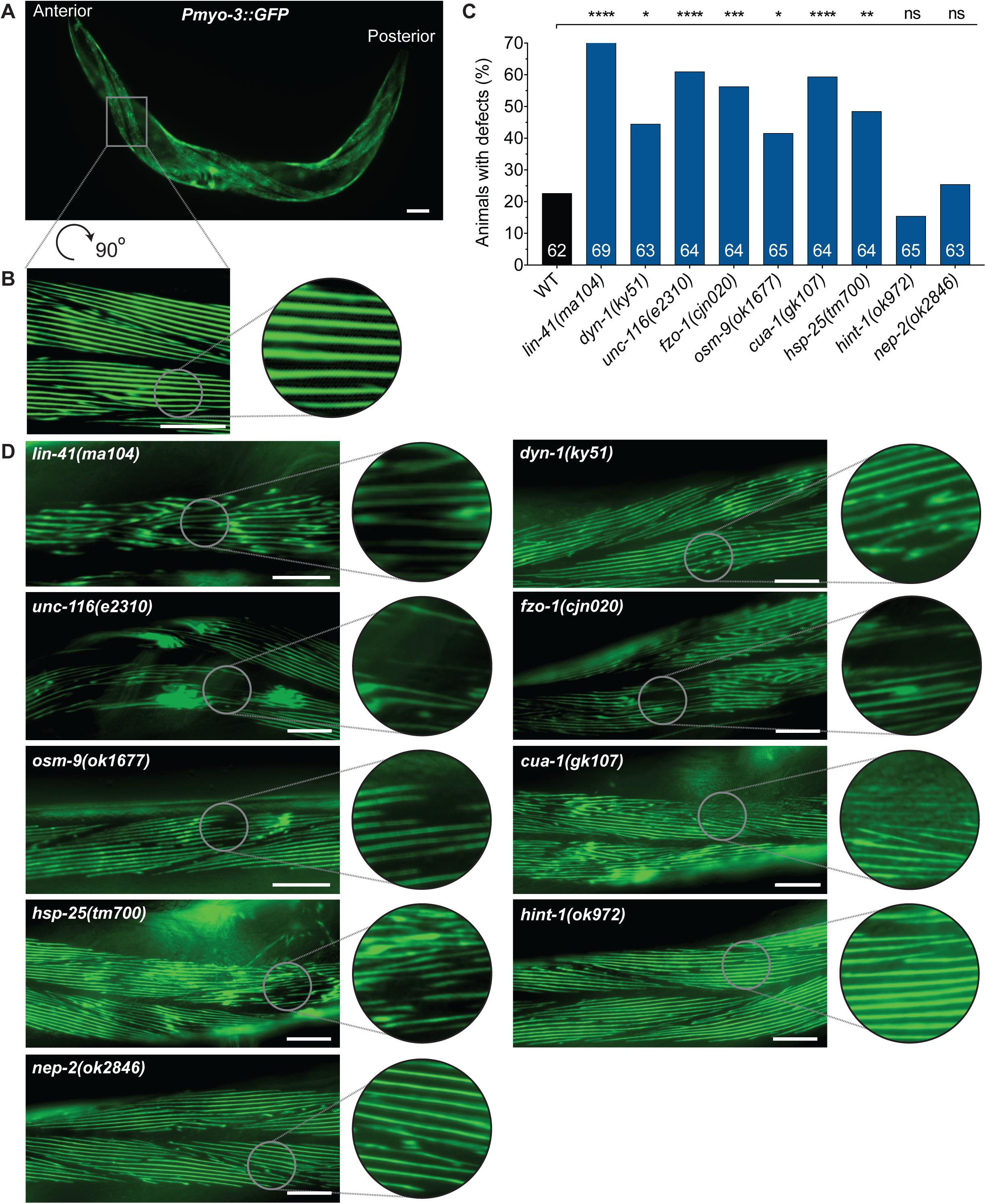
Visual scoring of body wall muscle defects. (**A**) A single 3-day old adult animal carrying the *stEx30*(*Pmyo-3::gfp::myo-3 + rol-6(su1006)*) transgene, which drives GFP-tagged myosin heavy chain in the body wall muscles. Animals with *stEx30* also possess a rolling phenotype that facilitates visualization of muscle cells that flank the dorsal and ventral sections of the body. (**B**) 90 ° rotated and expanded view of muscle area within dashed box in (A). (**C**) Categorical scoring of 3-day old adult WT and mutant animals, all carrying *stEx30*, for abnormalities in muscle fibres. Data is represented as the percentage of defective animals; *n* values are shown in each bar. Defects are compared between WT and CMT2 mutant animals using chi-square analysis with false discovery rate. \**P* < 0.05, \*\**P* < 0.01, \*\*\**P* < 0.001, \*\*\*\**P* < 0.0001, *ns* = not significant. (**D**) Representative close-up muscle fibre images of individual CMT2 mutant animals. Scale bar represents 60 µm in (A), and 25 µm in (B) and (D).

To understand the consequences of mutating the CMT2-related genes on muscle morphology, we used a transgene [*stEx30(Pmyo-3::GFP)*] to fluorescently label myosin heavy chain filaments within the body wall muscles (Fig. 4A, B). We assessed A3 stage worms for defects in filament integrity, scoring for loss of the distinct striations, GFP clumping due to accumulation of cellular debris, and the presence of gaps between the filaments as a sign of degeneration. Categorical scoring revealed that mutation of *lin-41* resulted in the highest proportion (∼70%) of defective animals, with most of the filaments in this group irregularly organized or degenerated with GFP aggregations (Fig. 4C, D). This was followed by *unc-116(e2310), cua-1(gk107)*, and *fzo-1(cjn020)* mutants, with greater than 55% of animals found to be defective in each of these backgrounds. The muscular organization appeared slightly better in these mutant backgrounds, although deterioration of filament lattices resulting in large gaps contributed to the high proportions of defects. GFP clumping was particularly prominent in *unc-116(e2310)* mutant animals. In comparison, the significant but smaller extent of muscle cell degeneration and deposits in *hsp-25(tm700), dyn-1(ky51)* and *osm-9(ok1677)* mutants meant that only 40-50% of animals were scored defective. Animals lacking *hint-1* or *nep-*2 were not significantly different from the wild-type, which surprisingly displayed structural defects in about 20% of animals. This may be due to aging-associated decline of muscle structure (Meissner et al., 2009).

With clear structural collapse of the myofilaments in the majority of mutant animals, we next sought to quantify these morphological changes in more detail. To the best of our knowledge, no protocols exist for quantitatively assessing morphological changes in *C. elegans* muscles, with qualitative scoring typically used to compare muscle defects. As such, we applied innovative but simple approaches with freely available tools to quantify two aspects of the body wall muscle phenotypes: cell area and myosin filament length. For the body wall muscle area, we calculated the total area of a single cell and the area of gaps within that selected cell using the polygon tool in Fiji software. The ratio of gap to total cell area was then calculated and compared between wild-type and mutant animals (Fig. 5A). In line with our categorical scoring results, the *lin-41(ma104)* mutant animals displayed the highest level of defects, with a 0.21 area ratio (Fig. 5B). This ratio was in stark contrast to that of wild-type, *hint-1(ok972)* and *nep-2(ok2846)* mutants, where the ratio was 10-fold lower at approximately 0.02. The ratio of gaps in the muscle cells of *unc-116, fzo-1, osm-9*, and *cua-1* mutants were also significantly increased compared to the wild-type (Fig. 5B). In contrast, and despite a considerably high number of *dyn-1(ky51)* and *hsp-25(tm700)* worms being scored as defective in our categorical scoring (Fig. 4C), the gaps within the muscle cells pertaining to structural deficits were inconsequential, resulting in no significant differences to the wild-type (Fig. 5B).

**Figure 5.**
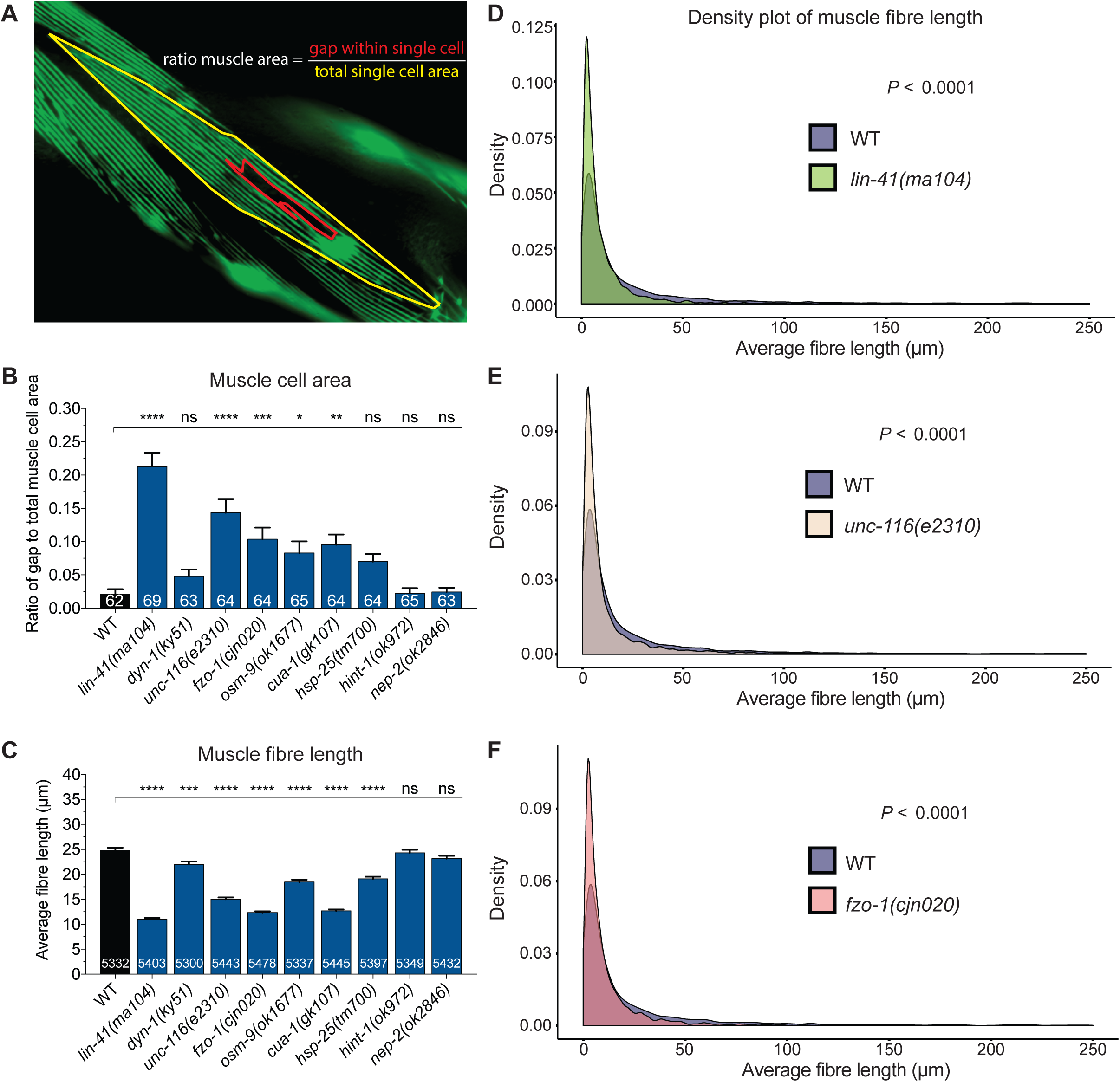
Quantification of body wall muscle area and fibre length in 3-day old adults. (**A**) Example image of how the muscle area ratio (gap to total cell area) was calculated. Gaps left by degenerating muscle fibres (red-lined area) and the total area of a single muscle cell (yellow-lined area) were drawn and calculated in Fiji using the polygon selection tool. (**B**) Comparison of gap to total muscle cell area ratio between WT and CMT2 mutants. (**C**) Measurement of individual fibre length using a combination of Fiji skeletonization and ilastik segmentation. The protocol is illustrated in Figure S3. Only images with at least one complete visible oblique muscle cell were included for analysis. Fibres that recorded 0 µm or longer than 250 µm were excluded. (**D-F**) Density plot of muscle fibre length compared between WT and *lin-41(ma104), unc-116(e2310)*, and *fzo-1(cjn020)*. Density plots of other CMT2 mutants can be found in Figure S4. Bar represents mean ± S.E.M in (B) and (C). Number of animals and fibres analyzed is listed in each bar in (B) and (C), respectively. \**P* < 0.05, \*\**P* < 0.01, \*\*\**P* < 0.001, \*\*\*\**P* < 0.0001, *ns* = not significant from one-way ANOVA with Dunnett’s post hoc tests for multiple comparison in (B) and (C). *F* test was used for variance comparison between WT and mutants in (D) to (F), with significance set at *P* < 0.05. All animals expressed the *stEx30* transgene.

Next, we used the ilastik image segmentation software to calculate the length of myofilaments (Fig. S3A). We exploited the ‘suggest features’ and ‘live update’ functions in the training window to help us determine if the labelling had been sufficient. Once satisfactorily segmented, the images were exported as PNG binary images, and opened in Fiji software (Fig. S3B). Default thresholding was then applied to the PNG image before skeletonization, a Fiji plugin, was implemented to filter out the border pixels (Fig. S3C). Following skeletonization, detailed analysis of the length of each filament was performed using the “Analyze Skeleton” plugin. Any fibre measurement of 0 µm was omitted, as this was not physiologically possible, and any measurement greater than 250 µm was also excluded as we calculated this to be the maximum reasonable length for a filament. Using this analysis, we observed significantly lower average fibre lengths in seven of the nine genetic mutations (Fig. 5C). In particular, the fibre lengths of *lin-41(ma104), fzo-1(cjn020)* and *cua-1(gk107)* worms were more than 50% shorter compared to wild-type, while *hint-1(ok972)* and *nep-2(ok2846)* worms were again unchanged. Mutations in *osm-9* and *hsp-25* led to a moderate 6 µm decrease in average filament length compared to wild-type, while the larger muscle gaps in *unc-116(e2310)* mutants further reduced the mean filament lengths by 10 µm. Despite the low ratio of gap to total muscle cell area in *dyn-1(ky51)* mutants, the average fibre length was significantly shortened by 3 µm in these animals (Fig. 5C).

We further analyzed the morphological differences in muscle structure by comparing the variances of filament length using density plots. In wild-type animals, the average filament length was more widely distributed compared to most of the mutants, with a higher proportion of longer filaments evident. In contrast, *lin-41(ma104), unc-116(e2310)* and *fzo-1(cjn020)* mutants had significantly smaller distributions of filament lengths (Fig. 5D-F). All of the remaining mutants except for *nep-2(ok2846)* also displayed smaller ranges of filament lengths (Fig. S4). In summary, we optimized quantitative methods to study the structural deficits in the body wall muscles, and found that mutations in genes associated with CMT2 led to increases in the ratio of gaps to total cell area, and considerably shorter myosin fibres. These methods eliminate potential bias typical of phenotypic scoring and introduce reliable means for quantifying muscle cell morphology. Moreover, they reveal muscle degeneration is a prominent feature associated with the mutation of CMT2-associated genes in *C. elegans*.

### Loss of CMT2-associated genes impacts muscle function

Next, we questioned if the morphological defects in the body wall muscles caused functional deficits. To specifically analyze the functionality of the body wall muscles, we exposed A3 stage worms to different concentrations of levamisole (40 µM and 200 µM). Levamisole is a potent agonist of nicotinic acetylcholine receptors, located postsynaptically on muscle cell membranes, and is commonly used to analyze the contractility of the body wall muscles. We measured the body length before and after levamisole treatment, as well as the time taken for the worms to completely paralyze. There were only negligible differences compared to the wild-type for most of the mutants in the relative change in body length (Fig. 6A), signifying minimal changes in muscle contraction and body stiffness. However, this was not the case for *osm-9(ok1677), cua-1(gk107)* and surprisingly, *nep-2(ok2846)*, which all demonstrated significantly reduced changes in body length in response to levamisole (Fig. 6A). Strikingly however, when we calculated the time taken for the animals to reach complete paralysis, all mutants except for *hsp-25(tm700)* and *hint-1(ok972)* were completely paralyzed by 40 µM in less than two-thirds of the time required by wild-type worms (Fig. 6B). The wild-type and mutant animals were all paralyzed more quickly when exposed to a higher concentration of levamisole, but *lin-41, dyn-1, unc-116*, and *fzo-1* mutants again displayed significantly faster responses. Curiously, the shorter time taken for some of the mutants to become fully paralyzed does not directly correlate with our results on locomotor impairment and the presence of structural defects within the body wall muscles. This indicates that muscle morphology and function are not strictly dependent, and may also indicate that the mutations affect nicotinic acetylcholine receptor signalling within the muscles in different ways, only some of which culminate in deficits in the specific behavioural paradigms we have tested. In summary, the phenotypic characterization and levamisole-induced contractions of body wall muscles show that mutations in CMT2 causal genes impair muscle morphology and function, contributing to decreases in mobility.

**Figure 6.**
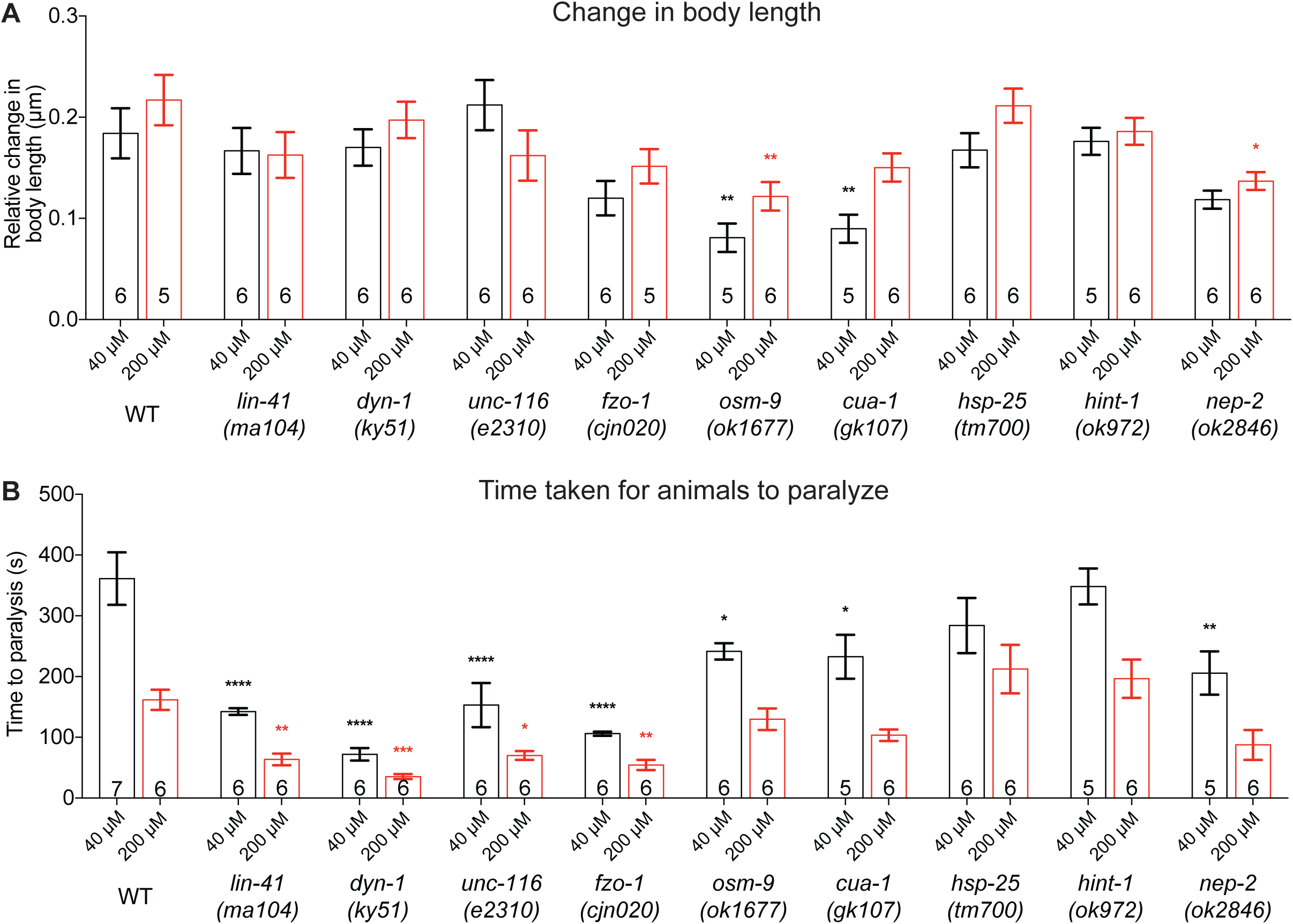
Quantification of body contraction in response to levamisole. (**A**) Relative change in body length of 3-day old adult wild-type (WT) and mutant animals following the application of different concentrations of levamisole (40 μM or 200 μM). The original and contracted body lengths were measured before levamisole administration and when the animals had been paralyzed by levamisole, respectively. (**B**) Bar graph of the time taken for the animals to reach full paralysis following the addition of 40 μM or 200 μM levamisole. Bars represent mean ± S.E.M, *n* values are within each bar. We used one-way ANOVA with Dunnett’s post hoc tests to compare WT and CMT2 mutant animals within each concentration. \**P* < 0.05, \*\**P* < 0.01, \*\*\**P* < 0.001, \*\*\*\**P* < 0.0001, *ns* = not significant.

### Electrophysiological recordings uncover neuromuscular junction defects

At the neuromuscular junction (NMJ), where axons innervate muscle cells, the axon terminals of motor neurons release acetylcholine to bind and activate postsynaptic nicotinic acetylcholine receptors to initiate muscle contraction. Compared to body wall muscles and neurons, the molecular function of NMJ is rarely studied in CMT animal models, even though loss of NMJ function has been reported in CMT2D mice (Chandhok and Soh, 2016, Spaulding et al., 2016). It has also been suggested that NMJ dysfunction is length-dependent because the distal extremities are more severely affected than proximal limbs in CMT2 patients (Saporta, 2015). Therefore, we next questioned if dysfunctional NMJ signalling may be a common deficit associated with mutating CMT2-associated genes, and whether it also contributed to the progressive decay in locomotion we observed in our *C. elegans* mutants. To analyze NMJ activity, we performed patch clamp electrophysiological recordings on the NMJs in the ventral nerve cord of A3 stage wild-type and CMT2 mutants. We focused on two properties of NMJ activity: the frequency and the amplitude of post-synaptic currents (PSCs). PSCs are generated when postsynaptic muscle receptors are activated by spontaneous acetylcholine release from presynaptic motor axon terminals. PSC frequency reflects the rate of spontaneous neurotransmitter release from presynaptic neurons while the amplitude is further dependent on postsynaptic membrane integrity and cholinergic receptor activity. Hence, optimal PSC frequency relies heavily on the activity of presynaptic motor neurons, whereas the functionality of both presynaptic motor neurons and postsynaptic body wall muscles could affect the amplitude of PSCs (Liu et al., 2013).

Dramatically, the frequency of spontaneous PSCs in *lin-41(ma104), dyn-1(ky51), unc-116(e2310)* and *fzo-1(cjn020)* animals was between 2- and 5-fold lower than the wild-type (Fig. 7A, C). The lower PSC frequencies of *unc-116(e2310)* and *dyn-1(ky51)* correlates with the morphological defects identified in the cholinergic motor neurons (Fig. 3C-E). Although we did not see a change in cholinergic motor neuron morphology as a result of the *lin-41* and *fzo-1* mutations, we did not analyze all cholinergic (or other classes of) motor neurons that may display degenerative phenotypes in these backgrounds. Nonetheless, the decrease in spontaneous PSC frequencies correlate with the potent deficits in both crawling and swimming, as well as the levamisole-induced muscle contractions, suggesting that presynaptic deficits likely play a major role in the movement defects and shorter response time to levamisole. In contrast, despite the numerous body wall muscle defects seen earlier, none of the mutant worms exhibited a significant decrease in mean amplitude of spontaneous PSCs (Fig. 7B, C). This could mean that the quantity, distribution and positioning of postsynaptic receptors are sufficient to offset the degeneration observed in some muscle cells. Nevertheless, our results provide strong evidence linking several genes implicated in CMT2 with robust NMJ signalling.

**Figure 7.**
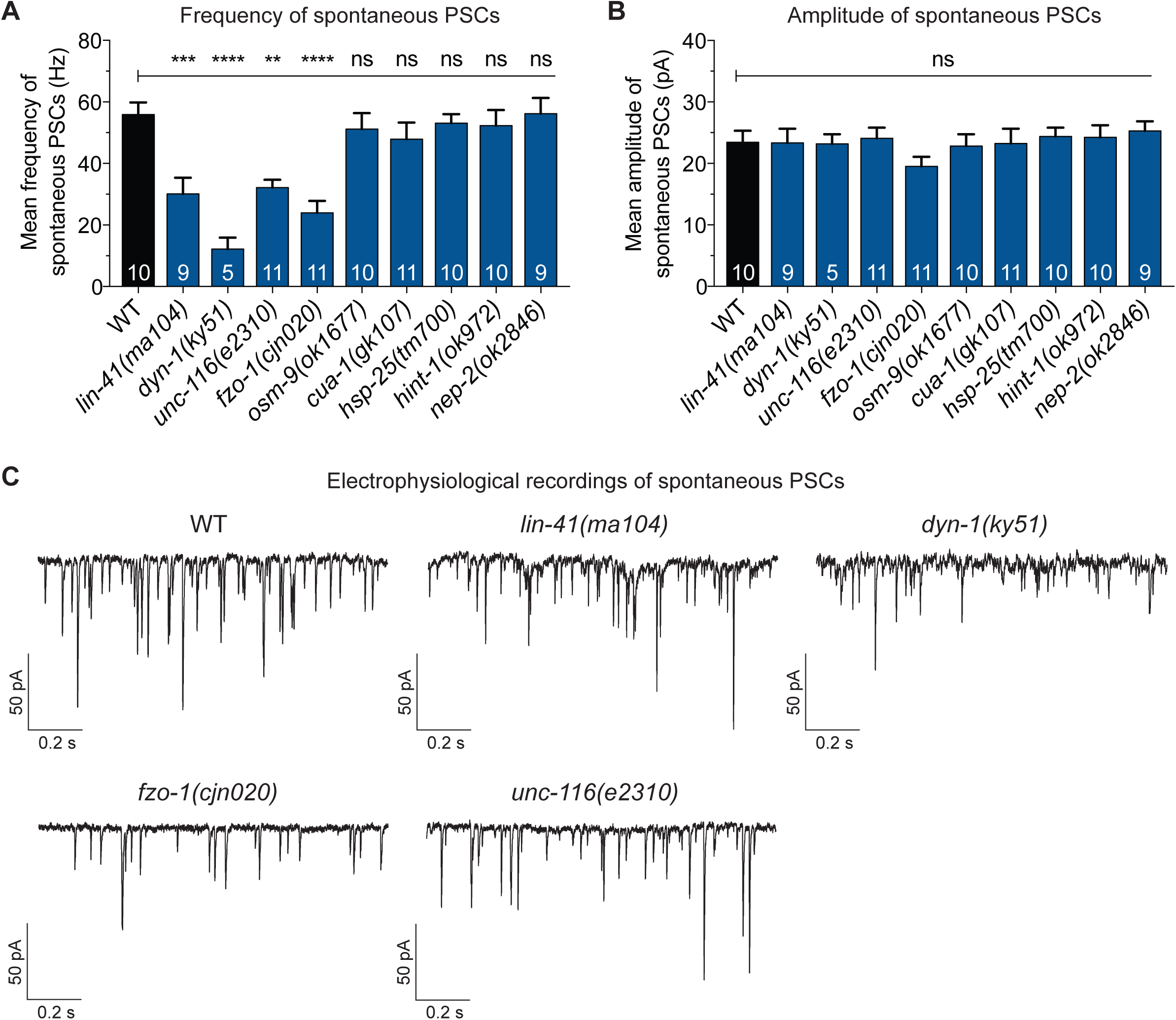
Electrophysiological recordings from the neuromuscular junction (NMJ). (**A**) Bar graph of the frequency of spontaneous postsynaptic currents (PSCs) recorded from 3-day old adult WT and CMT2 mutant animals. (**B**) Bar graph of the amplitude of spontaneous PSCs recorded from 3-day old adult WT and CMT2 mutant animals. (**C**) Sample traces of spontaneous PSCs recorded from ventral cord NMJs of individual worms. All recordings were made at a holding potential of –60 mV. All animals carried the *vsIs48* transgene. Bars represent mean ± S.E.M, *n* values are within each bar in (A) and (B). One-way ANOVA with Dunnett’s post hoc tests were used for comparison between WT and CMT2 mutant animals in (A) and (B). \*\**P* < 0.01, \*\*\**P* < 0.001, \*\*\*\**P* < 0.0001, *ns* = not significant.

## Discussion

Through the targeting of CMT2-related genes in *C. elegans*, our study identifies new insights into the importance of these genes for cellular function and animal behaviour. Overall, the nematode struggled with normal locomotion when these genes were mutated. Further investigations revealed that the locomotion defects were a consequence of structural breakdown of cholinergic motor neurons and degeneration of body wall muscles, as well as dysfunctional neuromuscular junction signalling. A summary of the main findings in this study is provided in Figure 8. Our results further underline the similarities between mammals and the nematode, revealing common behavioural, cellular and molecular traits between CMT2-associated mutants and CMT2 human patients. The development and characterization of *C. elegans* carrying mutations in CMT2-associated genes is an important development at a time when we have limited knowledge about the clinical and genetic heterogeneity of the disease and a complete lack of effective therapeutics.

**Figure 8.**
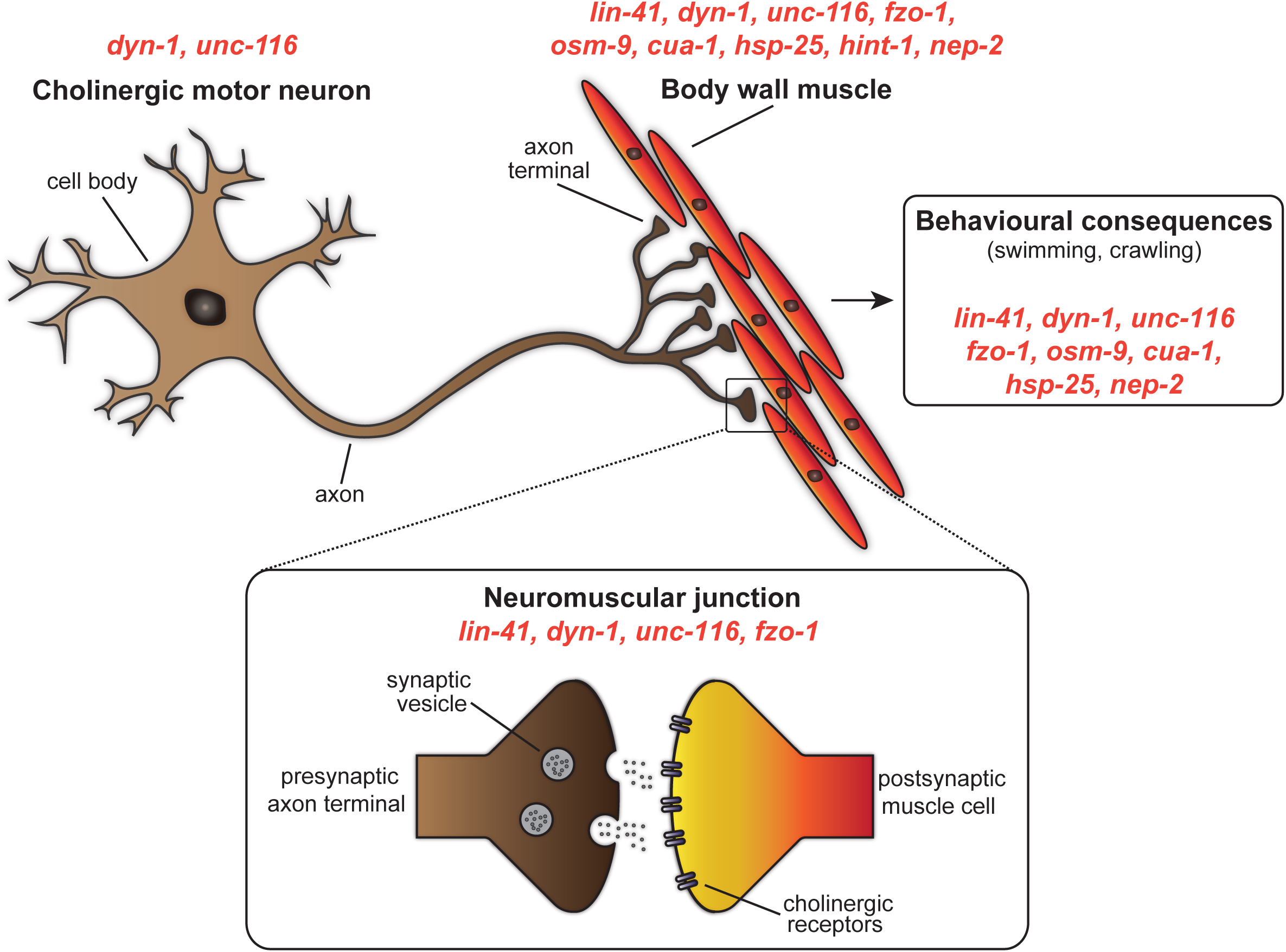
Summary of the sites at which mutation of CMT2-associated genes affect cellular function and animal behaviour in *C. elegans*. Mutations in *dyn-1* and *unc-116* led to defects in cholinergic motor neuron morphology, while mutations in *lin-41, dyn-1, unc-116, fzo-1, osm-9, cua-1, hsp-25, hint-1* and *nep-2* resulted in irregularities in the body wall muscle. Neuromuscular junction dysfunction occurred as a result of *lin-41, dyn-1, unc-116* and *fzo-1* mutations, whereas *lin-41, dyn-1, unc-116, fzo-1, osm-9, cua-1, hsp-25* and *nep-2* mutants experienced motility impairments in swimming and/or crawling.

### Shared characteristics between C. elegans and human patients following mutations in CMT2-associated genes

As CMT2 progresses, difficulty moving becomes the biggest obstacle for patients, mainly due to the loss of motor neurons, weakened muscles and limb deformities (Bird, 1993-2018). Locomotion analysis of the genetic mutants in our study revealed that *C. elegans* also experienced similar physical debilitation. In particular, worms carrying mutation in orthologous genes responsible for CMT2A (*fzo-1*/*MFN2*), CMT2C (*osm-9*/*TRPV4*), CMT2F (*hsp-25/HSPB1*), CMT2M (*dyn-1*/*DNM2*), CMT2R (*lin-41/TRIM2*) and unclassified-CMT2s (*unc-116*/*KIF5A, cua-1*/*ATP7A*) experienced declines in swimming and crawling ability as they aged. We speculated that these movement defects were a result of weakening body wall muscle and motor neuron structure, as these underlying impairments are also reported in CMT2 patients. Our hypothesis was first supported by the high proportion of DA/DB cholinergic motor neuron defects, especially in the absence of functional *unc-116* and *dyn-1* genes. The kinesin-1 heavy chain UNC-116/KIF5A is a microtubule-dependent motor protein expressed in neurons and muscle, which is required for the transport and localization of mitochondria and lysosomes, and for regulating neuronal polarity. Downregulation of kinesin-1 has been reported to reverse the axon-dendrite organization, leaving behind disordered neurites without distinct dendrites or axons (Yan et al., 2013). Similarly, DYN-1/dynamin-2 is a dynamin GTPase with overlapping cellular expression as UNC-116 that is involved in microtubule bundle processing and vesicular trafficking. *C. elegans* lacking functional DYN-1 displayed dysmorphic neuronal structure. However, human patients with mutations in dynamin-2 have presented contradicting neuronal features, with some studies reporting neuronal degeneration but not others. This could be due to the precise position of the mutations, with sequence changes within Pleckstrin homology domain and C-terminal proline-rich domain shown to induce more severe neuronal loss (Chandhok et al., 2017).

Our morphological analysis of body wall muscles revealed structural breakdown of myosin heavy chain filaments, again supporting the role of body wall muscle health on habitual locomotion. However, muscle biopsies are not often performed in patients, and accessible records show varied muscle descriptions among different subtypes of CMT2 (Ericson et al., 1998, González-Jamett et al., 2017, Renwick et al., 2016, Duis et al., 2016). Nevertheless, and consistent with our findings, many CMT2 patients carrying mutations in the orthologous genes for *fzo-1, unc-116, lin-41, hsp-25* and *osm-9* have been reported to exhibit muscular atrophy and myopathy especially in the distal limbs (Guinto et al., 2017, Landouré et al., 2010, Rouzier et al., 2012, Ylikallio et al., 2013, Houlden et al., 2008, Evgrafov et al., 2004, Barisic et al., 2007). As listed in Table S1, the functions of the genes and encoded proteins vary greatly between the subtypes, from outer mitochondrial membrane fusion (FZO-1) to osmotic sensory receptors (OSM-9) and ubiquitin ligases involved in gene regulation (LIN-41). Providing further complications, genes such as *cua-1*/*ATP7A* are also involved in other diseases. Mutation of *ATP7A*, which encodes a copper transporter, was identified as causative for CMT2 less than a decade ago (Kennerson et al., 2010), but has been associated with the neurodegenerative Menkes disease since 1993 (Vulpe et al., 1993). In hereditary neuropathy, missense mutations in *ATP7A* occur in a locus that results in distinct neuronal loss without causing the severe copper deficiency characteristic of Menkes disease(Kennerson et al., 2010). Interestingly, mutation of another gene with similar copper-binding functions, *SCO2*, was reported to cause a different type of axonal CMT with similar symptoms, thus highlighting the possible role of copper homeostatic imbalance in axonal neuropathy (Rebelo et al., 2018). Conversely, studies on CMT2M patients with dynamin-2 mutations mainly report normal muscle morphology, but occasional reports have described destabilization of skeletal muscle structure, further underlining the complexity of disease mechanisms (González-Jamett et al., 2017, Renwick et al., 2016). We also observed prominent GFP aggregation in the myosin filaments of some of the mutants, especially *unc-116*/*KIF5A*, which suggests underlying cellular toxicity, although the exact molecular mechanism behind this phenotype remains to be determined (Link et al., 2006, Meissner et al., 2009).

From the investigation of muscle function using levamisole, we found that some of the animals lacking CMT2-associated gene function experienced paralysis far more quickly than wild-type animals. This observation was somewhat unexpected, as the time taken to reach paralysis should be longer if the body wall muscles are impaired because less receptors would be available on damaged postsynaptic membranes for levamisole to bind. However, due to structural defects and presynaptic deficits, an increase in the number and sensitivity of postsynaptic receptors to agonists may occur as a compensatory mechanism for a reduced number of endogenous agonists at the NMJ. Hence, the addition and abundance of exogenous levamisole could activate the waiting postsynaptic receptors and thereby explain the shorter time taken by some of the mutants to paralyze.

Overall, our analysis of CMT2-associated genes in *C. elegans* revealed many common characteristics including difficulties moving, as well as functional declines in neuronal and muscle function. Further studies to precisely dissect the commonalities between the many causal genes and disease phenotypes will be important for a full understanding of their roles in CMT2.

### The need for viable CMT2 animal models

In order to better understand the underlying pathophysiology of CMT2, effective animal models are required. Unfortunately, the complexity of the disease has contributed to the scarcity of working animal models. Hence, numerous attempts to generate accurate animal models of CMT2, especially in rodents, have not been successful. For instance, constitutive *DNM2* knockout (*DNM2*^−/−^) is embryonic lethal in mice, while heterozygous *DNM2*^+−^ mice do not present any CMT2-like phenotypes (Zhao et al., 2018, Cowling et al., 2014). *DNM2*-deficient mice expressing a floxed DNM2 allele were found to be viable, but comprehensive analyses have not been performed, with only myelinated peripheral neurons analyzed (Sidiropoulos et al., 2012). Similarly, constitutive knockout in mice of *KIF5A*, the gene responsible for a number of neuromuscular disorders including CMT2, hereditary spastic paraplegia, and amyotrophic lateral sclerosis, results in lethality soon after birth. Karle et al. overcame this lethality with selective neuronal knockout of *KIF5A*, but insights into other cellular consequences such as muscle or NMJ phenotypes could not be made (Karle et al., 2012). Selective expression of pathogenic alleles in mouse neurons is also prevailing in CMT2A analysis, thus masking the effect of *MFN2* mutation on other cell types such as skeletal muscles. Mouse models have been generated for specific CMT2A mutations, including T105M and R94W, which have revealed important insights into the cellular mechanism of disease and therapeutic targets (Cartoni and Martinou, 2009, Feely et al., 2011, Bannerman et al., 2016, Strickland et al., 2014, Rocha et al., 2018). However, further complicating our understanding of the disease is the finding of different patient-specific mutations having alternative (and sometimes opposing) effects on protein function (Detmer and Chan, 2007, El Fissi et al., 2018). Furthermore, our findings on *HINT1* mutants largely mirrored those observed by Seburn and colleagues, where mice lacking functional HINT1 were phenotypically normal, indicating the need for a more effective animal model for *HINT1*-related disease (Seburn et al., 2014). Through detailed studies of muscle morphology, however, we could identify subtle changes in worms lacking *hint-1* that might impact behaviour in older animals or with additional genetic or environmental impacts. This highlights the need to appropriate animal models, as well as appropriate tools for identifying sites of pathology.

In addition to the unclassified CMT2s in this study, there have previously been no working animal models to explore the function of genes involved in CMT2C, CMT2R and CMT2T. In contrast to rodents, *C. elegans* knockouts are mostly viable, and mutation of the CMT2-associated genes in this study produced phenotypes similar to human patients, thus allowing us to properly study the effects of genes responsible for CMT2 on disease phenotypes in the nematode. In summary, this study of both behavioural and cellular consequences of mutating CMT2-associated genes in *C. elegans* is an important step forward in the establishment of reliable animal models to understand disease mechanisms. This is especially true for the unclassified CMT2 subtypes, as well as CMT2C, CMT2R and CMT2T, for which no animal models have been generated.

### C. elegans as an animal model for high-throughput drug screening

High-throughput screening (HTS) of small compounds on whole animals has been gaining momentum in recent years in bid to identify effective therapeutics that could improve disease symptoms. The main advantage of using whole animals in HTS is that the biological target does not need to be identified, as compounds can be screened based on phenotypic changes in motility or lifespan. This is a huge benefit for human diseases with poorly understood biological mechanisms such as CMT2. While rodents are highly favoured as disease models due to their closer similarities with humans, using rodents in HTS is not only extremely costly, but also time-consuming. In addition to the advantages associated with ease of culturing, small size, high progeny rate, and rapid life cycle, the *C. elegans* genome can be readily manipulated, and its transparent body makes visualization and analysis of cellular structures simpler than other animal models (Markaki and Tavernarakis, 2010). For these reasons, the nematode has been increasingly used as a tool for HTS.

One of the earliest large-scale, liquid-based compound screenings in *C. elegans* took place more than a decade ago, and focused on identifying novel antimicrobials and anti-ageing drugs (O’Reilly et al., 2013). Following the success of early HTS, a number of novel therapeutics that reversed the symptoms of a range of human diseases has since been identified in automated HTS assays using *C. elegans*. For instance, fluphenazine was discovered from HTS in *C. elegans* to be efficacious in reducing the accumulation of misfolded proteins responsible for α1-antitrypsin deficiency (ATS). The finding was subsequently replicated and validated in mammalian cell lines and a mouse model of the disease (Li et al., 2014). In a different study, Sleigh and colleagues modelled spinal muscular atrophy in *C. elegans* and combined the automated phenotyping system with HTS to identify three compounds, 4-aminopyridine (potassium channel blocker), gaboxadol (GABA_A_ receptor agonist), and N-acetylneuraminic acid (monosaccharide), that could improve motility in the mutant worms (Sleigh et al., 2011). More recently, similar restoration of movement by a number of drug classes (including neuroleptics) isolated from HTS in nematode models of amyotrophic lateral sclerosis and autism spectrum disorders was also discovered (Patten et al., 2017, Schmeisser et al., 2017). In the present study, we utilized the automated movement tracker WMicrotracker to detect the motility of worms in a high-throughput manner (96-wells per run). Consistent with manual and individual movement assays, a number of CMT2-asscoiated genetic mutants recorded a reduction in movement rate in the WMicrotracker. This phenotypic impairment has opened new doors of opportunity for high-throughput screening of small molecules on *C. elegans* CMT2 mutants, where compounds that successfully rescue the movement defect could be developed into effective therapeutics for the currently incurable disease.

### Beyond qualitative scoring: building the foundation for quantifying muscle morphology

The transparency of *C. elegans* allows for simple visualization and analysis of underlying cellular and organelle structures and processes. However, despite the large number of studies that have explored cellular defects as a result of genetic mutations, no reliable tools or measurements for quantifying changes in cellular morphology in *C. elegans* have been developed. Instead, categorical scoring has been the method of choice for defining defects. In 2014, Rizk and colleagues first optimized a “segmentation and quantification of subcellular shapes (SQUASSH)” protocol that enabled researchers to distinguish and quantify cellular and subcellular structures or shapes from fluorescence-based microscopy images (Rizk et al., 2014). Previously, we optimized the parameters in the SQUASSH segmentation ImageJ macro for a detailed, non-subjective assessment of mitochondrial morphology in both muscle and neuronal cells of *C. elegans* (Byrne et al., 2019). In this previous research, we also included movement assays, with the outcomes consistent with the current study, whereby animals carrying the *fzo-1(cjn020)* mutation experienced decreased movement compared to WT.

In the present study, we further improved on quantitative methods to non-subjectively analyze muscle defects. Using a combination of Fiji and ilastik segmentation software (Schindelin et al., 2012, Sommer et al., 2011), we were able to measure the muscle cell area and myosin fibre lengths for comparison with our qualitative scoring. From visual analysis of muscle cells, we saw conspicuous structural breakdown and irregularities of the myosin fibres especially in *lin-41, unc-116, fzo-1, osm-9* and *cua-1* mutants, which correlated with the lack of normal movement from these animals. Further examination of individual myosin fibre lengths and gaps using quantitative approaches also saw significant defects in the same mutants, thus validating our methodology. The lack of difference in the gap to total muscle cell area ratio and average fibre length between wild-type, *hint-1* and *nep-2* mutants was also consistent with the phenotypic scoring results, further supporting the viability of the quantitative measurements. Surprisingly, however, comparison of the relative density and variance of muscle fibre length revealed a sizeable difference between wild-type and all mutants except *nep-2*, although these differences may not necessarily lead to behavioural defects. Our results imply that comprehensive computational-based analyses can detect subtle changes in phenotypic variation that cannot be identified by visual means. Thus, by using Fiji and ilastik segmentation to quantify muscle cell morphology, we have taken a step forward in improving quantitative methodology to measure cellular phenotypic changes in *C. elegans*, and eliminated the potential bias accompanying categorical scoring.

### The neuromuscular junction as a target for therapeutics

To our knowledge, this study is the first to report a reduction in the frequency of spontaneous PSCs in *C. elegans* carrying mutations in genes associated with CMT2A, CMT2M, CMT2R, and the uncharacterized CMT2 caused by mutations in *unc-116*. Our findings suggest that these genes are important for robust NMJ function, and therefore identify this compartment as a potential therapeutic target for these disease subtypes. The presence of presynaptic cholinergic neuron defects and/or postsynaptic body wall muscle degeneration in these mutant animals further supports the notion of NMJ deficits, which are likely to further compound the motility impairments identified in the mutant animals. Deficits in NMJ function have previously been noted for other CMT classes, including CMT1A and CMT4 (Scurry et al., 2016, Cipriani et al., 2018, Ang et al., 2010). Furthermore, Spaulding, Sleigh and colleagues first demonstrated that synaptic defects could be a contributing factor to the early symptoms of axonal CMT (Sleigh et al., 2014, Spaulding et al., 2016). NMJ loss was observed in the proximal limb muscles in a CMT2D mouse model, which led to muscle weakness and restricted locomotion. It is also intriguing to note that administration of physostigmine, a reversible inhibitor of acetylcholinesterase which acts at the synapse, improved the movement of CMT2D mice, thus supporting the role of NMJ as a biological target for the development of CMT2 therapeutics (Spaulding et al., 2016). Further studies assessing the effects of physostigmine and other synaptically-acting drugs on NMJ function and motility in CMT2 animal models will be important for delineating the involvement of the NMJ in CMT2 pathophysiology. Taken together, the presence of NMJ abnormalities in animals lacking CMT2-asscoiated genes suggests that the NMJ could be a major site in the disease pathology, and additional studies looking into the properties of NMJ in the presence of CMT2 genetic mutations will provide a greater understanding of the role of NMJ stability in CMT2 pathophysiology and as a therapeutic target.

## Conclusions

We have shown that *C. elegans* carrying mutations in genes associated with CMT2 share many disease characteristics with human patients. This study has further highlighted the advantages of *C. elegans* as a reliable animal model to study neuropathies, and through the development of the first quantitative methods for analyzing muscle defects has laid the foundation for future comprehensive studies not just involving the *C. elegans* body wall muscles, but also potentially other cellular types. Our study of nine orthologous genes causing CMT2 has revealed common sites of deficiencies when these genes are lacking (Fig. 8), with functional loss at the NMJ and within the musculature most prominent. Our extensive characterization of CMT2-associated genetic mutants in *C. elegans* provides valuable insights and tools for understanding the common pathways underlying the heterogeneous neuropathy, and for future high-throughput drug screening approaches aimed at identifying novel therapeutics for the currently incurable and debilitating disease.

## Materials and Methods

### Generation and maintenance of C. elegans strains

Maintenance, crosses, and other genetic manipulations were all performed via standard procedures (Brenner, 1974). Hermaphrodites were used for all experiments and were grown at 20 °C on nematode growth medium (NGM) plates (0.25% peptone, 51 mM NaCl, 25 mM KH_2_PO_4_, 5 µg/ml cholesterol, 1 mM CaCl_2_, 1 mM MgSO_4_, 2% agar) seeded with OP50 *Escherichia coli*. A full list of strains used in this study is shown in Table S1.

### Swimming assays

Individual synchronized worms (L1 larvae stage, L4 larvae stage, 3-day old adult and 7-day old adult) were transferred to an unseeded 60 mm plate at room temperature to remove leftover OP50, then individual worms were transferred to a 10 μL droplet of M9 buffer (22 mM KH_2_PO_4_, 42 mM NaH_2_PO_4_, 86 mM NaCl, 1 mM MgSO_4_) (Brenner, 1974). Worms were left to acclimatize for 15 seconds, and the number of thrashes per minute counted using a hand counter under an Olympus SZ51 microscope. A total of 10 animals were used for each replicate, with three replicates performed. For analysis of swimming speed, videos of 5 worms per replicate swimming in 1 mL of M9 buffer on single unseeded 35 mm plate were recorded. These videos were then converted in Fiji (version 2.0.0) (Schindelin et al., 2012) to 30 fps frame rate in .*avi* format and swimming speed of animals within field of view was quantified using WormLab software (version 3.1.0, MBF Bioscience). The settings for swimming assays were adjusted according the protocol provided on the WormLab website (MBF Bioscience, 2018). Recordings were performed in triplicates or more, depending on the number of animals within the field of view. Data of any worm that went out of focus during recording was discarded. All images and videos were taken with Leica Microsystems M80/MC190 camera microscope. All assays were performed at room temperature.

### Body bends

Individual synchronized worms (L1 larvae stage, L4 larvae stage, 3-day old adult and 7-day old adult) were gently placed on unseeded 60 mm NGM plates and left to acclimatize for 3-5 minutes at room temperature. The number of body bends, defined as the maximum bend of the part of worm just behind the pharynx from one end to the opposite direction in a forward sinusoidal pattern, was manually counted for 3 minutes under the Olympus SZ51 microscope (Koelle, 2006). The reverse bend in the same direction was not included in the count. The number of body bends per minute was then averaged. To quantify crawling speed, videos of 5 worms per replicate crawling on a single unseeded 35 mm plate were recorded and converted in Fiji (version 2.0.0) (Schindelin et al., 2012) to 15 fps frame rate in .*avi* format. Crawling speed of the worms within field of view in the converted video was then quantified using WormLab software (version 3.1.0, MBF Bioscience). The settings for crawling assay were based on the protocol on the WormLab website (MBF Bioscience, 2018). Recordings were performed in triplicates or more. Data of any worm that went out of focus during the recording was discarded. All images and videos were taken with a Leica Microsystems M80/MC190 camera microscope. All assays were performed at room temperature.

### WMicrotracker

Large population of age-synchronized worms was prepared according to the Solis and Petrascheck protocol, with mild modifications (Fig. S2) (Solis and Petrascheck, 2011). Briefly, gravid hermaphrodites were bleached to release eggs, which were then hatched overnight at 20 °C to allow the newly-hatched worms to arrest at L1 larval stage. The L1 worms were plated the next day onto OP50-seeded 60 mm NGM plates and incubated at 20 °C until they reached adult stage. Each day until the day of experiment (3-day old adults), the worms were washed and separated from their progenies through gravitational precipitation onto new seeded plates. On the day of experiment, synchronized 3-day old worms were initially washed off the plates with M9 buffer into micro-centrifuge tubes, and subsequent washing of the worms in the tubes were performed at least twice to remove residual OP50 and offspring. 100 μL of 30-50 worms were then plated into each well of 96-well plate, 6 wells per genotype across two replicate experiments. The plate was then placed in the WMicrotracker-One™ instrument (PhylumTech) and analyzed for 3 hours at room temperature.

### Fluorescence imaging

The strains expressing the *stEx30*(*Pmyo-3::gfp::myo-3 + rol-6(su1006))* transgene were used for analysis of body wall muscles, while the *vsIs48*(*Punc-17::gfp*) transgene was used to visualize cholinergic neurons. Worms were immobilized with 0.05% tetramisole hydrochloride solution and mounted on 4% agarose pads on glass slides. Imaging of both body wall muscle and cholinergic neuron was carried out using Zeiss Axio Imager M2 microscope on 3-day old adult worms at 400 X magnification. For body wall muscle, the images were captured from the upper or lower part of the worm, excluding the extreme anterior and posterior regions, and regions adjacent to the vulva. For cholinergic neurons, only the dorsal neurons were analyzed. Both the defects of body wall muscle and cholinergic motor neurons were scored categorically as defective or not defective.

### Quantitative measurements of body wall muscle cell area and fibre length

To quantify body wall muscle cell area, we made sure that every image taken included at least one complete muscle cell (yellow enclosed region in Fig. 5A). Using the polygon selection in Fiji (version 2.0.0), we measured the area size of a muscle cell and the gaps within each cell. The ratio of the gaps relative to the total muscle cell area was then calculated. For this analysis, 60 or more worms were imaged and analyzed. For muscle fibre (myosin filament) length, 5 to 11 body wall muscle images (*.tif* format converted from *.czi* in Fiji version 2.0.0) of each mutant strain were first segmented in ilastik (version 1.3.0) (Sommer et al., 2011) (Fig. S3). During segmentation, the individual muscle fibres were carefully “trained” or classified to separate them from the unwanted background. Each segmented image was then exported (output data type unassigned 8-bit, range renormalized from 0,1 to 0,255) as a viewable binary image in *.png* format. The *.png* files were next threshold-adjusted (default setting) and skeletonized in Fiji (version 2.0.0, Skeletonize 2D/3D plugin). The lengths of the ‘skeletons’, or individual myosin filaments, were then analyzed using the Fiji Analyze Skeleton 2D/3D plugin (Fig. S3C). Filaments that recorded 0 or more than 250 µm (the maximum reasonable length for a filament) were excluded. Muscle cells with sections that were out of focus were also not included in the analysis.

### Quantification of muscle contraction using levamisole

To quantify levamisole-induced body contraction, animals were first incubated in 10 μL drug-free solution (140 mM NaCl, 5 mM KCl, 5 mM CaCl_2_, 5 mM MgCl_2_, 11 mM dextrose, and 5 mM HEPES; 330 mOsm, pH adjusted to 7.2) to measure the original body length. This was followed by measurement of body length after the addition of 10 μL of 80 μM or 400 μM levamisole solution to obtain a final concentration 40 μM or 200 μM levamisole. The time taken for the worms to reach full paralysis after being incubated in levamisole solution was recorded. Worm body was monitored under Olympus MVX10 microscope equipped with XM10 CCD camera. The cellSens software (version 1.7, Olympus) was used to calculate the length of the worm body.

### Electrophysiology

Whole-cell patch-clamp recordings of NMJs were performed on 3-day old adult animals at room temperature under a 60 X water immersion lens with an EPC-10 amplifier and the Patchmaster software (HEKA) (Richmond and Jorgensen, 1999, Liu et al., 2013). Voltages were clamped at −60 mV. Recordings of spontaneous PSCs from each worm were analyzed for the frequency and amplitude of spontaneous PSCs using MiniAnalysis (Synaptosoft, Inc.). Recording pipettes (4–6 MΩ) were pulled from borosilicate glass and fire polished. The pipette solution contained 120 mM KCl, 20 mM KOH, 4 mM MgCl_2_, 10 mM HEPES, 0.25 mM CaCl_2_, 36 mM sucrose, 5 mM EGTA, and 4 mM Na_2_ATP (315 mOsm; pH adjusted to 7.2). The bath solution contained 140 mM NaCl, 5 mM KCl, 5 mM CaCl_2_, 5 mM MgCl_2_, 11 mM dextrose, and 5 mM HEPES (330 mOsm; pH adjusted to 7.2).

### Statistical analysis

Statistical analysis and graphs were performed/generated using Prism 7 (GraphPad Software) or R version 3.2.2 (R Core Team) (R Core Team, 2014). One-way ANOVA was used for comparing means of more than two groups followed by Dunnett’s multiple comparison post hoc test. *F* test was used to compare variances. Chi-square test (*chisq.post.hoc* R package), controlled for false discovery rate, was used to compare categorical data. Significance was set at *P* < 0.05 unless stated otherwise.

## Acknowledgements

We thank Joe Byrne for generating the *fzo-1(cjn020)* strain; Marina Kennerson and Ebru Boslem for comments on the manuscript, and members of the Neumann lab for valuable discussions and input. Some strains were provided by the CGC, which is funded by NIH Office of Research Infrastructure Programs (P40 OD010440). The authors thank WormBase for its wealth of information on *C. elegans*, and acknowledge Monash Micro Imaging, Monash University, for the provision of instrumentation, training and technical support. We are also grateful to Ruidong Xiang for his help and advice on R programming. This work was supported by CMTAA research grants (2015 and 2018), and NHMRC Project Grant 1099690 awarded to BN.

## Authors’ contribution

MS and BN designed the study. MS performed and analyzed all of the experiments and data except for the levamisole assays performed by XC and the electrophysiological recordings conducted by JL. MS and BN wrote and edited the manuscript. All of the authors read and approved the final manuscript.

---

**Figure S1. Movement assays across various *C. elegans* life stages**. (**A**) Thrash rates compared between wild-type (WT) and the nine mutant strains, across three different ages, larval stage 1 (L1), larval stage 4 (L4), 3-day old adults (A3) and 7-day old adults (A7). (**B**) Rate of body bends of WT and CMT2 mutant animals quantified across the same ages as in (A). Each dot in (A) and (B) represents a single animal (*n* ≥ 30). One-way ANOVA with Dunnett’s post hoc tests were used to compare rates of thrash or body bend between WT and mutant animals in (A) and (B). Data is represented as mean ± S.E.M. \*\**P* < 0.01, \*\*\**P* < 0.001, \*\*\*\**P* < 0.0001, *ns* = not significant.

**Figure S2. Age-synchronizing populations of *C. elegans*.** Schematic of the workflow, from synchronization via bleaching and washing, to plating of synchronized 3-day old adult animals in a 96-well plate for experimentation.

**Figure S3. Body wall muscle segmentation protocol using Fiji and ilastik.** (**A**) Original muscle segment must contain at least one full visible oblique muscle cell. (**B**) Distinct myosin fibre is classified and segregated from the unwanted background (in asterisk) in ilastik. (**C**) Skeletonization of image in Fiji. The image was skeletonized to filter out the border pixels, leaving behind only the skeletal remnants that become the topological representations of the original fibres. Measurement of each fibre was performed following skeletonization. Fibres that were 0 µm or more than 250 µm were excluded. Scale bar represents 25 µm.

**Figure S4. Density plot of muscle fibre length for each genotype.** (**A-F**) The variance of all except for *nep-2(ok2846)* was statistically different compared to wild-type (WT). Statistics were performed using *F* test for variances, significance set at *P* ≤ 0.05. Experiments were performed on 3-day old adult animals that carried *stEx30* transgene.

**Table S1.** List of strains used in this study.

