## Supplementary material for "Disruption of genes associated with Charcot-Marie-Tooth type 2 lead to common behavioural, cellular and molecular defects in *Caenorhabditis elegans*": Table S1

**Supplementary Table 1**. List of strains used in this study.

| Strain name | Gene (allele) | Type of mutation | Number of times outcrossed | Coded protein (function) |
|---|---|---|---|---|
| RW1596 | *myo-3(st386); stEx30(Pmyo-3::gfp::myo-3 + rol-6(su1006))* | Not annotated<br><br>GFP-tagged rescue | - | Myosin heavy chain |
| LX929 | *vsIs48(Punc-17::gfp)* | GFP-tagged WT | - | - |
| BXN418 | *lin-41(ma104)* | Transposon insertion (Slack et al., 2000) | 3 | RBCC Ubiquitin Ligase (protein degradation, gene regulation) |
| BXN674 | *lin-41(ma104); vsIs48(Punc-17::gfp)* | | | |
| BXN679 | *lin-41(ma104); myo-3(st386); stEx30(Pmyo-3::gfp)* | | | |
| BXN567 | *dyn-1(ky51)* | Missense substitution (P70S) (Clark et al., 1997) | 3 | Dynamin GTPase (endocytosis, synaptic vesicle recycling, cytokinesis, degradation of apoptotic cells) |
| BXN678 | *dyn-1(ky51); vsIs48(Punc-17::gfp)* | | | |
| BXN685 | *dyn-1(ky51); myo-3(st386); stEx30(Pmyo-3::gfp)* | | | |
| FF41 | *unc-116(e2310)* | Not curated | 15 | Kinesin 1 Heavy Chain (transport and localisation of synaptic vesicle components, axonal transport of neurofilament proteins) |
| BXN675 | *unc-116(e2310); vsIs48(Punc-17::gfp)* | | | |
| BXN680 | *unc-116(e2310); myo-3(st386); stEx30(Pmyo-3::gfp)* | | | |
| BXN248 | *fzo-1(cjn020); zdIs5* | 2629 bp deletion (Byrne et al., 2019) | 3 | GTPase (outer mitochondrial membrane fusion) |
| BXN623 | *fzo-1(cjn020); vsIs48(Punc-17::gfp)* | | | |
| BXN366 | *fzo-1(cjn020); zdIs5(Pmec-4::GFP); myo-3(st386); stEx30(Pmyo-3::gfp)* | | | |
| BXN377 | *osm-9(ok1677)* | 1478 bp deletion (Deletion Mutant Consortium, 2012) | 3 | TRPV channel (osmo- and warmth sensor channel) |
| BXN625 | *osm-9(ok1677); vsIs48(Punc-17::gfp)* | | | |
| BXN628 | *osm-9(ok1677); myo-3(st386);* | | | |

| | | | | |
|---|---|---|---|---|
| | *stEx30(Pmyo-3::gfp)* | | | |
| BXN562 | *cua-1(gk107)* | 1639 bp deletion (Deletion Mutant Consortium, 2012) | 3 | Copper-transporting P-type ATPase (copper transporter) |
| BXN626 | *cua-1(gk107); vsIs48(Punc-17::gfp)* | | | |
| BXN629 | *cua-1(gk107); myo-3(st386); stEx30(Pmyo-3::gfp)* | | | |
| BXN620 | *hsp-25(tm700)* | 870 bp deletion (Deletion Mutant Consortium, 2012) | 3 | Heat Shock Protein (molecular chaperone, actin organisation, axonal transport of neurofilament proteins) |
| BXN676 | *hsp-25(tm700); vsIs48(Punc-17::gfp)* | | | |
| BXN681 | *hsp-25(tm700); myo-3(st386); stEx30(Pmyo-3::gfp)* | | | |
| BXN619 | *hint-1(ok972)* | 1025 bp deletion (Deletion Mutant Consortium, 2012) | 3 | Histidine Triad Nucleotide Binding Protein 1 (nucleotide hydrolytic activity) |
| BXN624 | *hint-1(ok972); vsIs48(Punc-17::gfp)* | | | |
| BXN627 | *hint-1(ok972); myo-3(st386); stEx30(Pmyo-3::gfp)* | | | |
| BXN542 | *nep-2(ok2846)* | 362 bp deletion (Deletion Mutant Consortium, 2012) | 3 | Neprilysin (signalling peptide regulation) |
| BXN677 | *nep-2(ok2846); vsIs48(Punc-17::gfp)* | | | |
| BXN683 | *nep-2(ok2846); myo-3(st386); stEx30(Pmyo-3::gfp)* | | | |


BYRNE, J. J., SOH, M. S., CHANDHOK, G., VIJAYARAGHAVAN, T., TEOH, J.-S., CRAWFORD, S., COBHAM, A. E., BORALESSA YAPA, N. M., MIRTH, C. K. & NEUMANN, B. 2019. Disruption of mitochondrial dynamics affects behaviour and lifespan in *Caenorhabditis elegans*. *Cellular and Molecular Life Sciences*, 1-19.

CLARK, S. G., SHURLAND, D. L., MEYEROWITZ, E. M., BARGMANN, C. I. & VAN DER BLIEK, A. M. 1997. A dynamin GTPase mutation causes a rapid and reversible temperature-inducible locomotion defect in *C. elegans*. *Proc Natl Acad Sci U S A,* 94**,** 10438-43.

DELETION MUTANT CONSORTIUM 2012. Large-scale screening for targeted knockouts in the *Caenorhabditis elegans* genome. *G3 (Bethesda),* 2**,** 1415-25.

SLACK, F. J., BASSON, M., LIU, Z., AMBROS, V., HORVITZ, H. R. & RUVKUN, G. 2000. The *lin-41* RBCC gene acts in the *C. elegans* heterochronic pathway between the *let-7* regulatory RNA and the LIN-29 transcription factor. *Mol Cell,* 5**,** 659-69.
