## Supplementary figures and images for "Disruption of genes associated with Charcot-Marie-Tooth type 2 lead to common behavioural, cellular and molecular defects in *Caenorhabditis elegans*"

### Figure S1

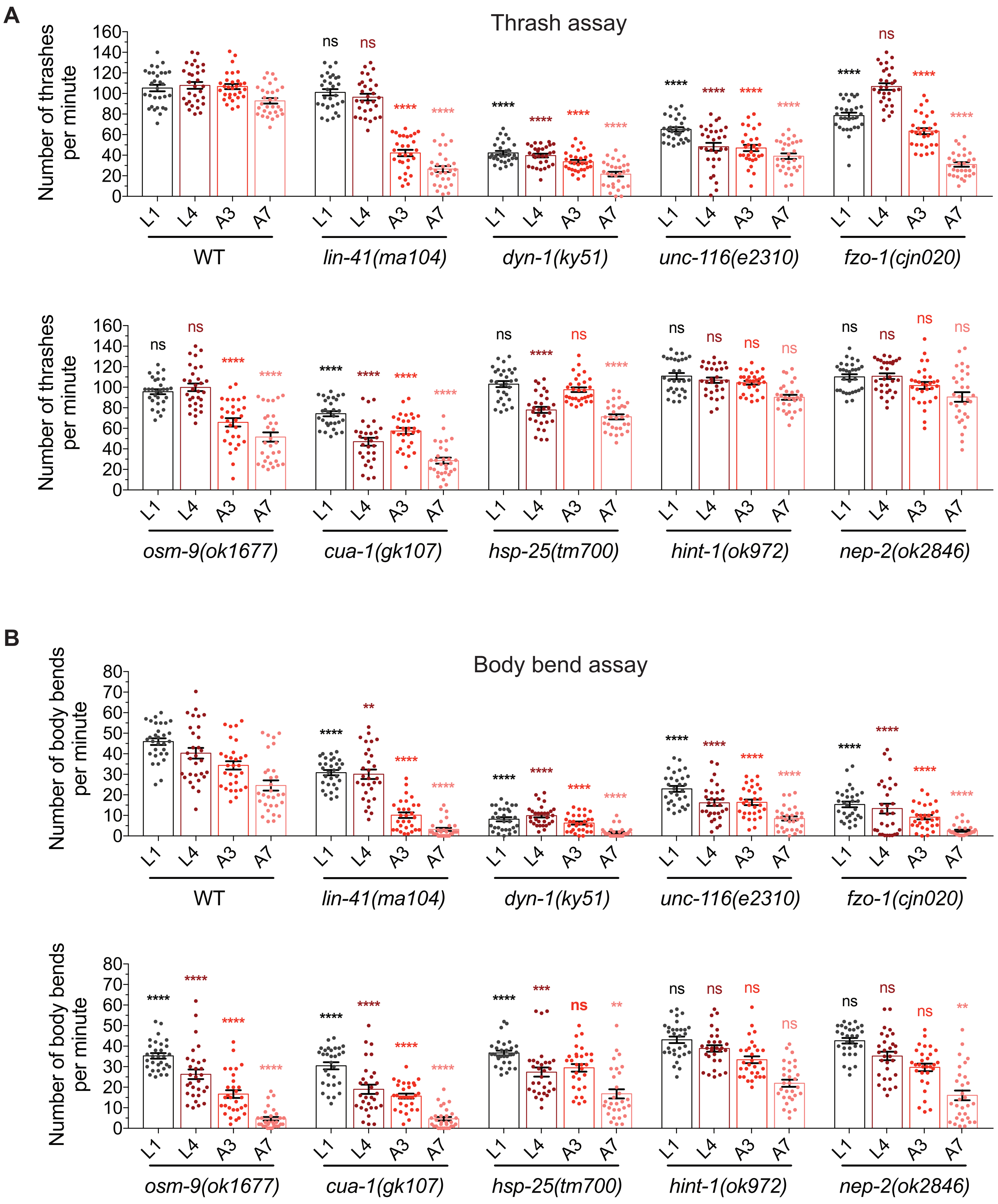

### Figure S2

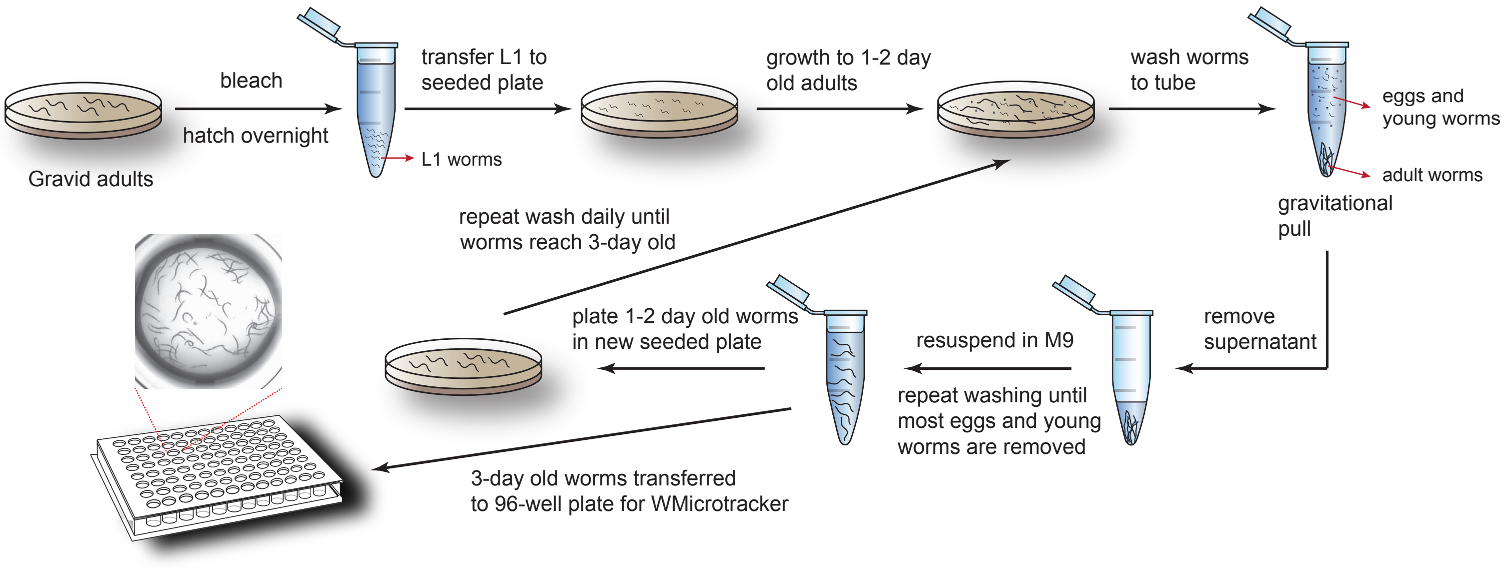

### Figure S3

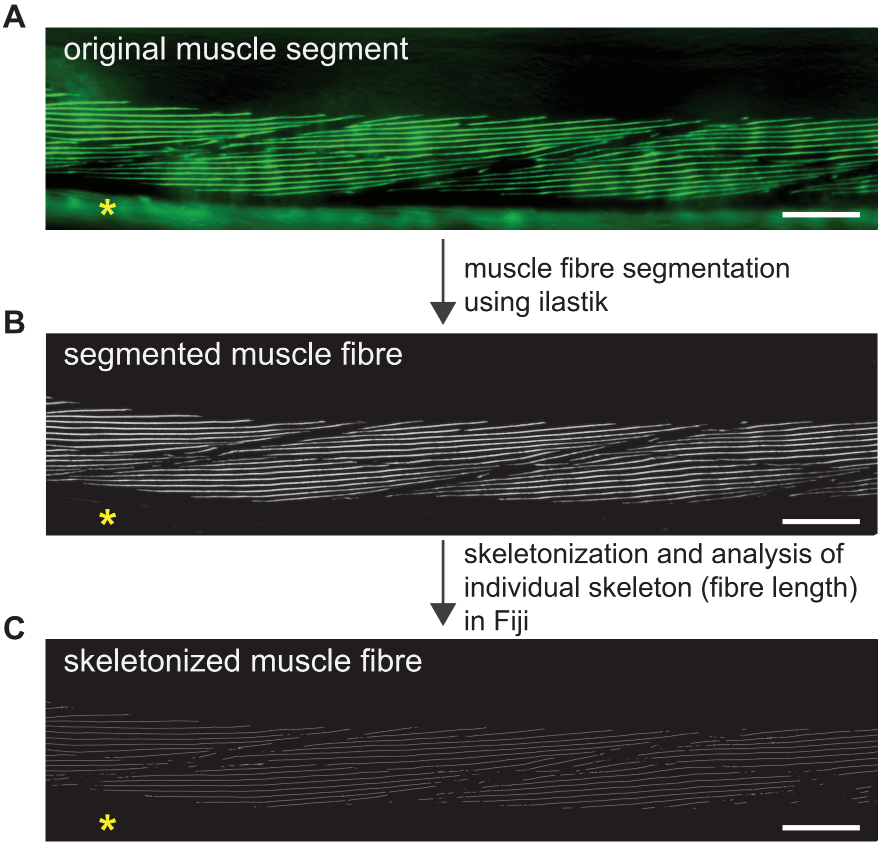

### Figure S4

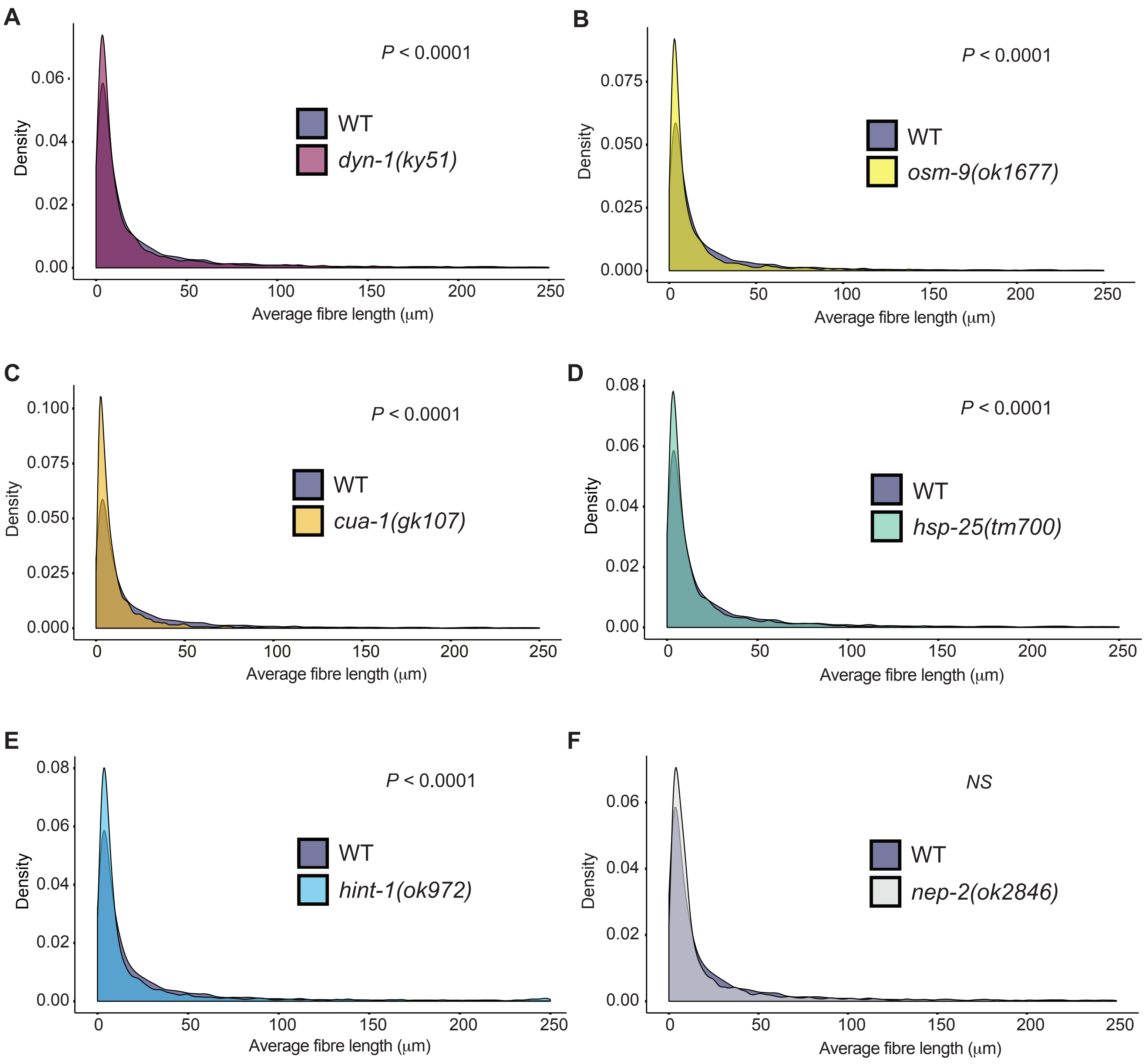
